# Revisiting the number of self-incompatibility alleles in finite populations: from old models to new results

**DOI:** 10.1101/2021.12.20.473438

**Authors:** Peter Czuppon, Sylvain Billiard

## Abstract

Under gametophytic self-incompatibility (GSI), plants are heterozygous at the self-incompatibility locus (*S*-locus) and can only be fertilized by pollen with a different allele at that locus. The last century has seen a heated debate about the correct way of modeling the allele diversity in a GSI population that was never formally resolved. Starting from an individual-based model, we derive the deterministic dynamics as proposed by Fisher (1958), and compute the stationary *S*-allele frequency distribution. We find that the stationary distribution proposed by Wright (1964) is close to our theoretical prediction, in line with earlier numerical confirmation. Additionally, we approximate the invasion probability of a new *S*-allele, which scales inversely with the number of resident *S*-alleles. Lastly, we use the stationary allele frequency distribution to estimate the population size of a plant population from an empirically obtained allele frequency spectrum, which complements the existing estimator of the number of *S*-alleles. Our expression of the stationary distribution resolves the long-standing debate about the correct approximation of the number of *S*-alleles and paves the way to new statistical developments for the estimation of the plant population size based on *S*-allele frequencies.

## 1 Introduction

Gametophytic self-incompatibility (GSI) is a genetically controlled mating system, common in flowering plants. GSI prevents crossing between individuals sharing identical alleles at the self-incompatibility locus (*S*-locus), especially self-fertilization (Durand et al., 2020). Despite extensive empirical and theoretical studies from the seminal works by Emerson (1938) and Wright (1939), many properties of the evolutionary dynamics of GSI are not well understood yet. For instance, the stationary distribution of *S*-allele frequencies or the expected number of *S*-alleles in a finite population are still not clearly established. Yet, such predictions are necessary to interpret genetic data in GSI populations. In addition, the probability of invasion of a mutant, or an estimator of effective population size from genetic data, both classical quantities in population genetics, are still lacking. By using modern stochastic tools, we address these questions in this manuscript.

It was early observed that *S*-allele diversity in natural populations was surprisingly large: in a population of *Oenothera organensis* at least 45 different *S*-alleles had been identified despite the population comprising at most one thousand individuals (Emerson, 1938, 1939; Lewis, 1949b). The large S-allele diversity is now known to be prevalent in GSI populations across various species and families (Castric and Vekemans, 2004). In the 1930’s, explaining the origin, dynamics and maintenance of this diversity was a real challenge for the young fields of population and evolutionary genetics. Wright (1939) was the first to propose an approximation of the stationary distribution of *S*-allele frequencies in a finite population, one of the first applications of theoretical population genetics as introduced by Fisher (1930) and Wright (1937). Wright (1939) used his approximation of the stationary distribution to estimate the expected number of *S*-alleles in a given population of finite size with recurrent mutation. His prediction, applied to *O. organensis* populations, was much lower than the observed number of *S*-alleles (Emerson, 1938) under a reasonable mutation rate and population size. Wright (1939) then suggested that the discrepancy between theoretical prediction and empirical observation could be due to population subdivision of the *O. organensis* population.

The failure of Wright’s attempt to apply a population genetic model to an empirical case generated a fierce and long debate about the good formulation of stochastic models of population genetic dynamics. Fisher (1958) and Moran (1962) were the first to criticize Wright’s initial model, mostly for a lack of mathematical rigor. Wright further refined his GSI model (Wright, 1960, 1964), and provided, with others, computer simulations (Kimura and Crow, 1964; Ewens and Ewens, 1966; Mayo, 1966) that confirmed that his approximation of the stationary distribution of *S*-allele frequencies, as well as his prediction of the expected number of *S*-alleles, were correct.

Yet, to the best of our knowledge, there is still no theoretical derivation and justification of the allele frequency dynamics and stationary state in a GSI population, even though, more than 50 years ago, Moran (1962, p.163) already suggested that the probabilistic model underlying GSI should be properly specified. Wright (1964) tentatively answered Moran’s criticism by framing it as “a basic difference in viewpoint”. Ewens and Ewens (1966) then argued that “[Wright’s] approximations are far better than could reasonably be expected”. An explicit derivation of the (macroscopic) properties of a GSI population from the individual (microscopic) level is necessary to determine the assumptions under which the approximation remains valid, and therefore can be used to interpret empirical observations and to conduct parameter inference. Indeed, paraphrasing Ewens and Ewens (1966): the validity of Wright’s approximation is “possibly fortuitous”, as suggested by some of the results by Mayo (1966) where the approximation can be incorrect (see Ewens (1964) and Ewens and Ewens (1966) for technical arguments).

In the end, one question remains unsolved: Wright’s approximation of the GSI stationary distribution seems to work well, but we do not know why. Here, we address this theoretical gap. We start by defining an individual-based model of gametophytic self-incompatibility. Taking the infinite population size limit, we first obtain the deterministic dynamics of gametophytic self-incompatibility, an equation that has already been derived and studied in great detail (Fisher, 1958; Nagylaki, 1975; Boucher, 1993). Second, we derive a diffusion approximation and compute the stationary distribution based on the central limit theorem for density-dependent Markov processes (Kurtz, 1971). We obtain an explicit expression for the stationary distribution that only depends on the population size and the number of *S*-alleles in the population, which formally confirms the validity of Wright’s approximation under general conditions. Finally, we also provide new results: 1) the invasion probability of a novel *S*-allele in a resident population, and 2) a method to estimate the effective population size from *S*-allele frequency data, based on our explicit result of the stationary distribution of *S*-allele frequencies.

## 2 Stochastic model of gametophytic self-incompatibility

We consider a diploid plant population with fixed size *N* with overlapping generations (Moran-type model). The analogous model definition of the self-incompatibility dynamics with non-overlapping generations (Wright-Fisher-type model) is given in the Supplementary Material (SM), Section C. We assume a fixed number of *S*-alleles in the population and denote it by *M*. The number of plants with genotype {*i j*} is denoted *A_i j_* with no allele order (*i.e. A_i j_* = *A_j i_*). Because of GSI, a {*i j*} individual can only be fertilized by pollen of type *k* ≠ *i*, *j* and then produces an offspring of type {*i k*} or {*j k*} with equal probability, consequently *A_i i_* = 0 for all *i*. The dynamics of the number of {*i j*} individuals is described in terms of coupled birth and death events to maintain the fixed population size *N*, as is common in Moran-type models. Then, the number of {*i j*} individuals, *A_i j_*, increases by one if

(b-i) a type *i* pollen fertilizes a {*j*•} individual that transmits allele *j*, or
(b-ii) a type *j* pollen fertilizes a *{i*•} individual that transmits allele *i*,

and a non-{*i j*} individual dies and is replaced.

The probability of fertilization of a {*jk*} individual with an *i* pollen is given by *p_i_*/(1 – *p_j_* – *p_k_*), where *p_i_* is the frequency of *i*-type pollen in the population 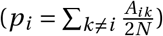. The form emerges from conditioning on successful fertilization, which reflects the assumption that the number of pollen is essentially infinite, or at least not limiting the (female) reproductive success of a plant, *i.e*. we assume no pollen limitation.

The rate at which the number of {*i j*} individuals increases by 1, the *birth rate*, is then given by:

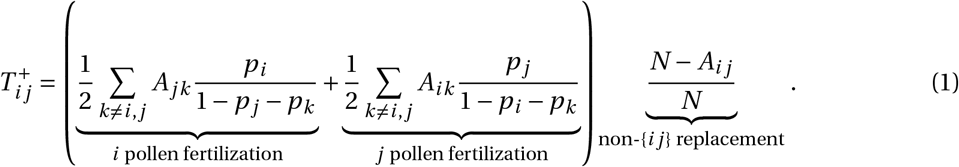

The reproduction terms between brackets were first derived by Fisher (1958).

Using the same arguments, the number of {*i j*} individuals decreases by one if

1. (d-i) a non-{*i j*} plant is fertilized (by any pollen), or
2. (d-ii) a *{i*•} plant is fertilized but the offspring is not {*i j*}, or
3. (d-iii) a {*j*•} plant is fertilized but the offspring is not {*i j*},

and a {*i j*} individual dies and is replaced.

The rate at which the number of {*i j*} individuals decreases by one, the *death rate*, is then given by (detailed derivation in the SM, Section A):

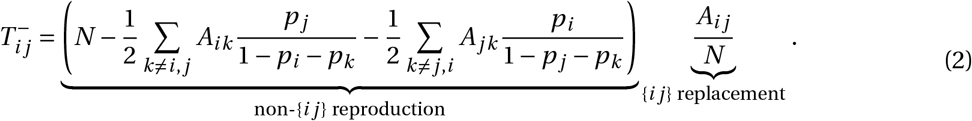

Since all individuals are equally likely to reproduce (at rate 1), *i.e*. the overall rate of reproduction is *N*, the reproduction part of the death rate is given by *N* minus the reproduction rate of an {*i j*} individual (Eq. (1)), which explains the term between brackets in Eq. (2).

Last, there is also the possibility that no change in the number of *i j* individuals occurs. This is the case if an *i j*-individual is produced and replaces an *i j* individual or a non-*i j* individual is produced and replaces a non-*i j* individual. The rate for this to happen is 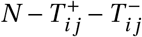.

We now proceed with the theoretical analysis of this stochastic model.

## 3 Results

### 3.1 Approximation of the dynamics in a finite population

First, we derive an approximation of the dynamics of genotypes and *S*-alleles in a large but finite GSI plant population. We assume that the number of *S*-alleles in the population is constant, *i.e*., we ignore the generation of new *S*-alleles through mutations and the loss of *S*-alleles through stochastic extinction events. To derive the genotype and *S*-allele dynamics, we apply diffusion-theoretical arguments (reviewed in the context of evolutionary applications in Czuppon and Traulsen (2021)). The mathematical details are stated in the SM, Section A. Briefly, if we denote by *a_ij_*(*t*) = *A_i j/N_* the proportion of {*i j*} individuals in the population at time *t*, we find an equation of the form

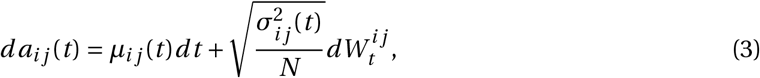

where *μ_i j_*(*t*) is the deterministic dynamics of the genotype frequencies (see below), 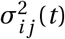 is the infinitesimal variance of genotype frequencies (explicitly given in SM, Section A, Eq. (A.4)), and 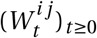 are Brownian motions, *i.e*. 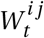 is normally distributed with mean zero and variance *t*, related to the stochastic fluctuations of genotype frequencies, which model the inherent randomness of births and deaths in a finite population. Notably the stochastic fluctuations scale with 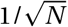 and vanish in the limit of infinite populations (*N* → ∞), where only the deterministic dynamics *μ_i j_* remain. The deterministic change of the genotype frequencies is given by

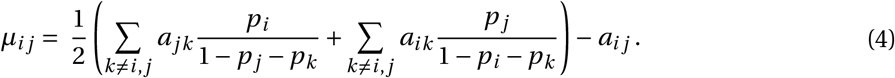

Solving *μ_i j_* = 0 in Eq. (4) gives the steady state 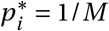 and 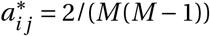 where all genotypes on the one hand, and all *S*-alleles on the other hand, have identical frequencies (Nagylaki, 1975; Boucher, 1993).

The diffusion approximation of the dynamics given in Eq. (3) describes genotype trajectories. The stochastic component of the equation, the second term on the right hand side, quantifies how strongly the genotype frequency fluctuates around the deterministic trajectory, the first term on the right hand side. These random fluctuations are of order 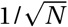, which means that they decrease with increasing population size *N* and vanish in the limit *N* → ∞ (we refer to Ethier and Kurtz (1986) for a rigorous treatment). The precise scaling of these stochastic fluctuations, which are a result of the finite population size, is obtained by explicitly computing the infinitesimal variance 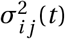 (SM, Section A).

The derivation of the *S*-allele dynamics is obtained by similar arguments and using the relation between the frequency of *S*-allele *i* and the genotype {*i j*} frequencies: *p_i_* = ∑*j*≠*i a_ij_*/2 (details in SM, Section A). The deterministic change in *S*-allele frequency, *i.e*. the infinitesimal mean, is then given by

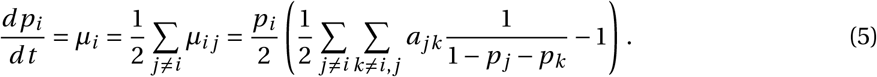

Eq. (5) shows that, since we assume no pollen limitation, the deterministic dynamics of allele *i* is driven by pollination of non-*i* plants (first term in the brackets), rather than pollination of *i*-plants by pollen of a different type (Fisher, 1958; Nagylaki, 1975; Boucher, 1993).

The diffusion approximation of *S*-allele frequencies is

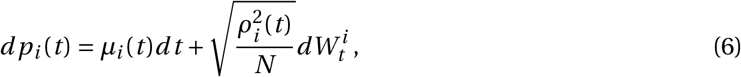

where 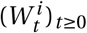 is a Brownian motion that reflects the stochastic fluctuations of the *i*-th *S*-allele frequency around the deterministic trajectory *μ_i_*. The exact expression of the infinitesimal variance of the *S*-allele frequency change 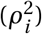 is computed in SM, Section A, Eq. (A.9). Note that the infinitesimal variance of *S*-alleles depends on the covariance between genotype frequencies because of our restriction of a fixed population size. Therefore, an increase in frequency of one *S*-allele (birth event) inevitably entails a decrease of another *S*-allele (death event).

### 3.2 Approximation of the stationary distribution

In an infinite plant population, the genotype and *S*-allele frequencies will converge to the steady states 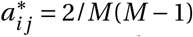 and 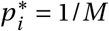, respectively. This deterministic steady state is globally stable (Boucher, 1993). This means that in a finite population the stochastic trajectories of *S*-allele frequencies will fluctuate around the value 1/*M*, at least as long as the probability of extinction of a *S*-allele is negligibly small. This will typically be the case if 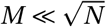. Using arguments motivated by the central limit theorem of density-dependent Markov processes (Ethier and Kurtz, 1986; van Kampen, 2007), we can then approximate the fluctuations around the deterministic steady state by a normal distribution (details in SM, Section B). We find that the stationary distribution of *S*-allele frequencies, denoted by *ψ*, is distributed as (denoted by the “~” sign)

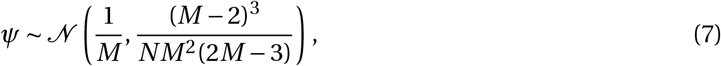

where 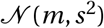 denotes a normal distribution with mean *m* and variance *s^2^*.

The same derivation is possible for a model with non-overlapping generations. In this case, we find that the stationary distribution differs only by a factor 2 in the denominator of the variance (SM, Section C, Eq. (C.9)), in line with the generally found difference between the variances of the Moran and Wright-Fisher model (Czuppon and Traulsen, 2021).

Fig. 1 shows that the approximation of the stationary distribution in Eq. (7) predicts well the stationary distributions obtained by stochastic simulations, especially if the number of *S*-alleles (*M*) is small. The approximation of the stationary distribution is slightly worse for large numbers of *S*-alleles because the probability that an allele is lost by chance increases with the number of *S*-alleles in the population, as shown by the bar located at zero for *M* = 15 in Fig. 1. As a consequence of an allele extinction, the mean allele frequency is displaced from 1/*M* to 1/(*M* – 1) in some simulations, or even lower values of *M* if multiple extinction events occurred. This explains the slightly worse fit of the approximated stationary distribution for *M* = 15, compared to lower numbers of *S*-alleles.

**Figure 1:**
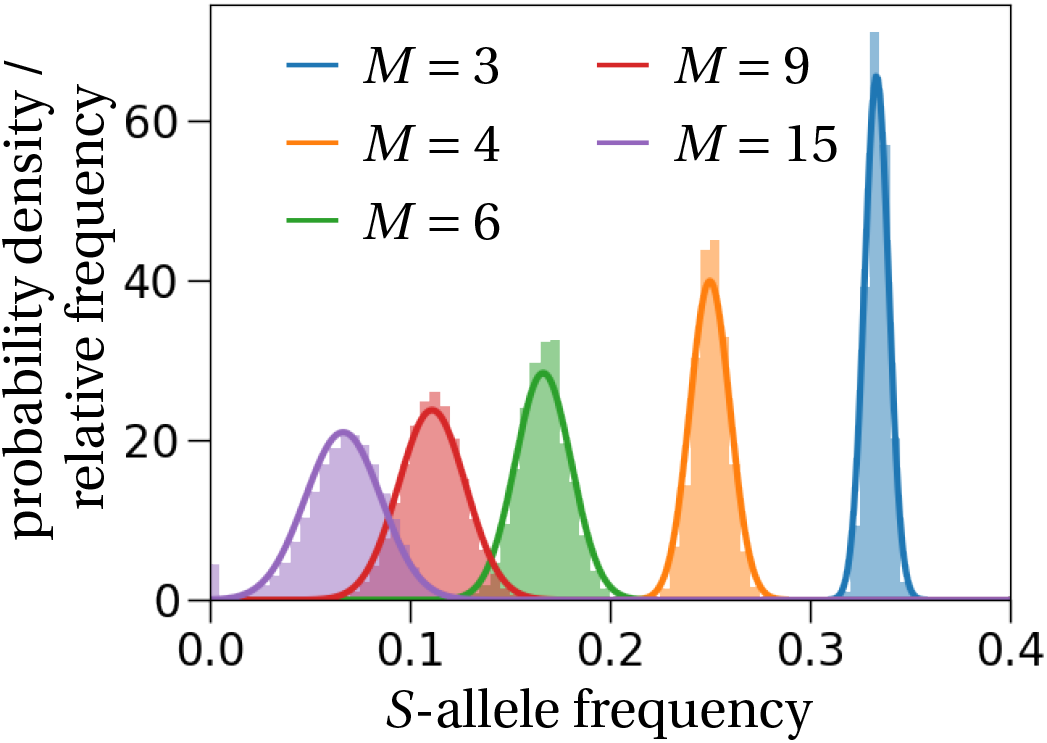
Stationary distribution of *S*-alleles. As predicted (solid lines, Eq. (7)), the stationary distribution of the allele frequency is well approximated by a normal distribution centered at 1/*M*. The histograms are calculated from 100 stochastic simulations that ran for 1,200 generations in a population of size *N* = 1,000, where we recorded the frequency of the same *S*-allele every generation. To ensure no dependence on the initial state we started recording frequencies after 200 generations. If the number of *S*-alleles becomes too large, *e.g. M =* 15, alleles might get lost, which is shown by the purple bar at the frequency equal to zero.

### 3.3 The number of S-alleles maintained in a finite population

We now allow the number of *S*-alleles to vary over time. In particular, at each reproduction event a mutation occurs and a new *S*-allele arises. The mutation probability is denoted by *u*. We assume an infinitely many alleles model, which means that every mutation will generate a new *S*-allele. Then, the rate at which new *S*-alleles arise is simply given by the mutation probability *u* (because the overall rescaled reproduction rate in the population is 1 due to the conditioning on successful reproduction).

To determine the number of *S*-alleles that are stably maintained in a population of finite size, we follow the partially heuristic reasoning by Wright (1939) and fill in the analytical gaps using the stationary distribution and the individual-based model. The basic idea is as follows (Wright, 1939): First, given an approximation of the stationary distribution of the allele frequencies for a given number of *S*-alleles *M* in a finite population of size *N* (Eq. (7)), the *loss rate* of a focal *S*-allele is computed by multiplying the probability for this *S*-allele to be at frequency 1/2*N* with its death rate *T*^−^ at frequency *1/2N* (computation in SM, Section D). Second, the loss rate is balanced with the *gain rate* of new *S*-alleles, which is just the mutation probability *u*. This yields the following equality

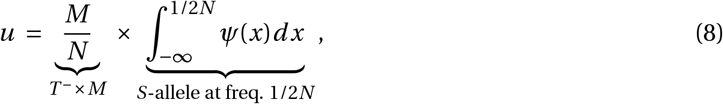

which can be solved numerically for *M*. Fig. 2 shows a comparison between the prediction obtained by solving Eq. (8) for different values of *M* and stochastic simulations. The number of *S*-alleles maintained in a finite population is reasonably well predicted by our model, even though it is overestimated as population size increases. The reason for this is that we underestimate the extinction risk of an *S*-allele in large populations. This is a consequence of our approximation of the stationary allele frequency distribution being a less accurate description of the dynamical system for large population sizes and a large number of *S*-alleles. The more *S*-alleles are found in a population, the less strong is the deterministic force due to self-incompatibility and the system dynamics appear more neutral. This explains why our derivation of the stationary allele frequency distribution, which is based on the assumption of a (strongly) attracting fixed point, becomes less accurate. This in turn leads to an overestimation of the number of stably maintained *S*-alleles in large populations.

**Figure 2:**
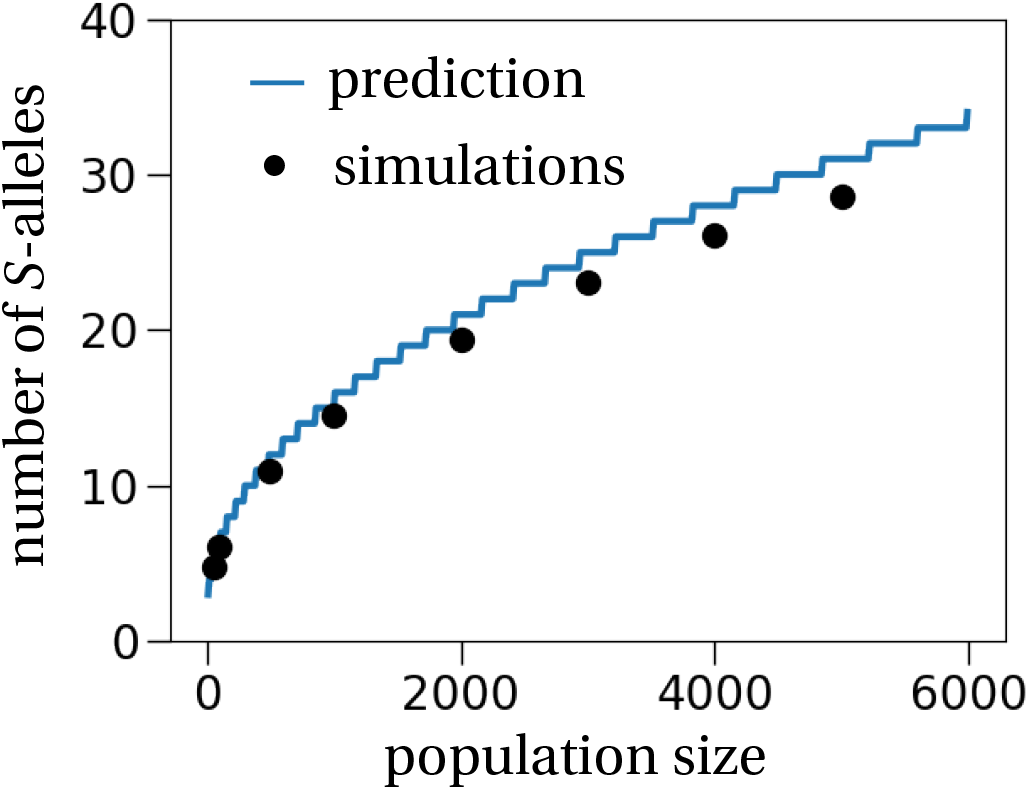
The number of *S*-alleles in a finite population. The general method to estimate the number of *S*-alleles as proposed by Wright (1939) and made explicit by our theoretical approach (solid line computed by Eq. (8)) predicts well the number of *S*-alleles in a finite population, obtained from stochastic simulations (dots). Each simulation was initiated with 30 different *S*-alleles and the number of *S*-alleles was recorded every 100 generations for 10,000 generations after a 10,000 generation burn-in period. The mutation rate was set to *u* = 1/(100*N*), *i.e*., one mutation every 100 generations on average. This procedure was repeated 50 times for every population size.

The overall good fit confirms the previous conclusion from numerical simulations (Ewens and Ewens, 1966; Mayo, 1966) that Wright’s (1939) general methodology indeed correctly predicts the number of *S*-alleles in a population. Note that we adapted Wright’s (1939) methodology for predicting the number of *S*-alleles in a finite population with two small adjustments: we used our new approximation of the stationary distribution (Eq. (7)) that is theoretically justified by the central limit theorem (yet neglecting allele covariances), and we used the explicit expression of the death rate of a single individual with the focal *S*-allele derived from the individual-based description of the model (instead of different intuitively proposed rates by Wright (1939, 1964)).

### 3.4 Invasion probability of a new *S*-allele

We now derive new results on the invasion behavior of a novel *S*-allele. In this section, we again ignore mutations and study the situation where at time 0, there is a single *S*-allele that is present in exactly one individual, *i.e*. this *S*-allele is at frequency 1/2*N*. All the other *S*-alleles are assumed to be in equilibrium, a reasonable assumption in view of the stability of the steady state (Boucher, 1993). The invasion probability can then be computed from the diffusion approximation (Eq. (6)) by applying results from stochastic diffusion theory (mathematical details in SM, Section E). We find that a new *S*-allele, appearing at frequency 1/2*N* by mutation (or immigration), establishes in a population with *M* resident *S*-alleles with probability

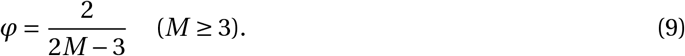

This approximation agrees well with simulation results (Fig. 3). Surprisingly though, it fits the simulation results better than the non-approximated formula from which Eq. (9) is derived; more details in SM, Section E.

**Figure 3:**
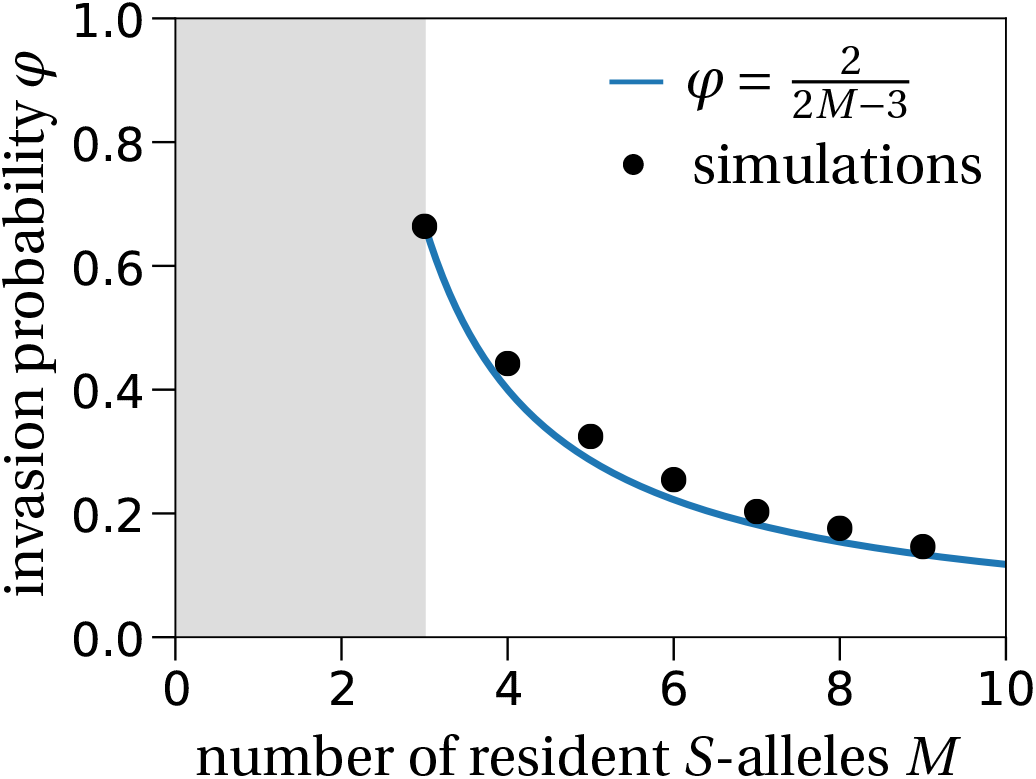
Invasion probability of a new *S*-allele. The invasion probability for different numbers of resident *S*-alleles (*M*) fits the result obtained from the diffusion approximation in Eq. (9) (solid line). The population size is set to *N* = 10,000 and the number of independent stochastic simulations is 10,000 per number of resident *S*-alleles. The grey region corresponds to the region where the plant population is not viable because at least three *S*-alleles are needed for successful reproduction.

The invasion probability decreases with *M* because the rare allele advantage is smaller when the number of resident *S*-alleles *M* increases. This is explained by an increase in the number of plants that can be fertilized by a single resident *S*-allele for larger numbers of resident alleles. The invading allele is initially present in a single copy, which means that the number of individuals it can fertilize is always *N* – 1. All the resident *S*-alleles, assuming that genotype frequencies are in equilibrium at 2/*M*(*M* – 1) (except for one genotype that is at frequency 2/*M*(*M* – 1) – 1/*N* because of the invading allele), can fertilize

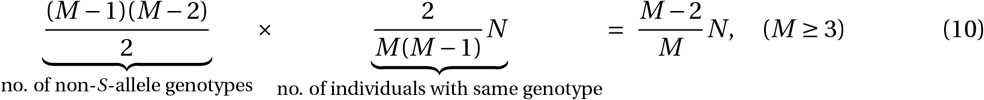

individuals. This shows that as *M* increases, the difference in the rate of successful pollen fertilization between the invading *S*-allele, *i.e. N* – 1, and the resident alleles, *i.e*. *N*(*M* – 2)/*M*, decreases. In contrast, since we assume no pollen limitation, the probability for a plant to be fertilized only depends on its genotype frequency in the population.

### 3.5 Estimation of the number of *S*-alleles and the population size from data

Our goal in this section is to provide a joint estimator of both the population size and the number of *S*-alleles in a sampled plant population genotyped at the *S*-locus. The main idea is to use the mean and the variance of the stationary distribution (Eq. (7)) to estimate the two population parameters of interest, *M* and *N*, given the sample size *n* and the number of *S*-alleles observed in the sample. We have developed two different estimators. The first estimator accounts for the sampling process and uses this correction to estimate the number of *S*-alleles and the population size. The second estimator computes the first two moments of the sample allele frequency distribution. These estimated moments are then used to estimate the mean and variance of the theoretical allele frequency distribution, which is then translated to the number of *S*-alleles and the population size.

The first estimator is an extension of Paxman’s (1963) urn model and the ideas presented in Yokoyama and Hetherington (1982) (details are given in SM, Section F). We derived a least-squares estimator for the number of *S*-alleles and the population size from the mean and variance of the number of occurrences *Y* of a given *S*-allele observed in the sample (SM, Section G). We show that the mean **E**[*Y*] and the variance **V**[*Y*] can be used to estimate the number of *S*-alleles and the population size. However, the minimization of the least-squares function showed convergence instability, presumably because the least-squares function is very flat along the axis of the population size (SM, Section G, Fig. C): Even though the number of *S*-alleles was correctly estimated, the population size estimator performed rather poorly (Fig. **4A**).

**Figure 4:**
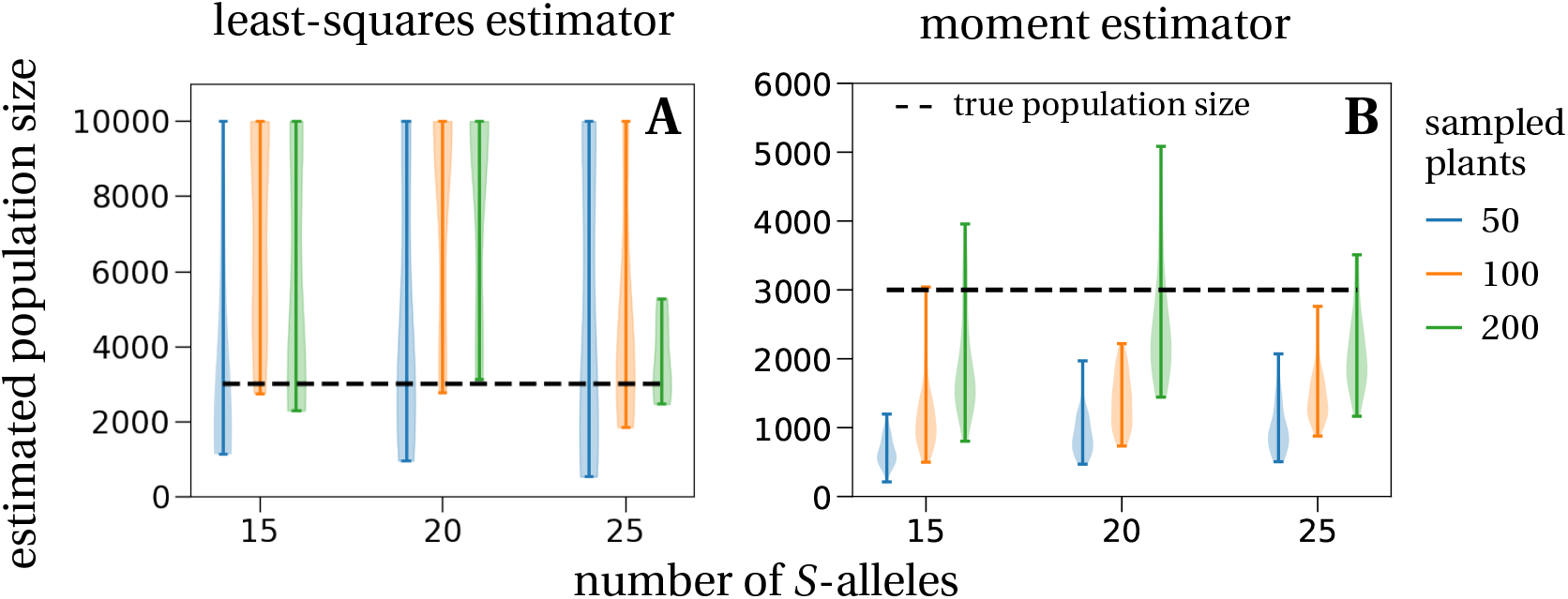
Estimated population sizes from simulated data. **A:** Estimated population sizes with the (weighted) least-squares method (details in Section G of the SM, Eq. (G.8)), which accounts for the sampling process, are widespread and mostly overestimating the true population size of 3000 plants (dashed line). This is true for all investigated numbers of *S*-alleles in the population (x-axis: 15, 20, 25) and all sample sizes (differently colored violinplots). The maximal allowed estimate is a population size of 10,000 plants. **B:** The second estimator, which is a moment estimator of the stationary allele frequency distribution (Eq. (11)), shows a tendency to underestimate the true population size (3,000 plants, dashed line; note the change of y-axis scope). This effect is most pronounced for small sample sizes (50 sampled plants, blue violinplots) and becomes less prominent for intermediate (100 sampled plants, orange violinplots) and large sample sizes (200 sampled plants, green violinplots). The violinplots are population size estimates for **A:** 10 and **B:** 50 different random samples taken from the same data set, implemented by a hypergeometric distribution (sampling without replacement). The simulated population of size 3,000 with 15, 20 or 25 different *S*-alleles was started close to its steady state.

The second estimator is a moment estimator and ignores the sampling process associated with data collection. We fit the theoretically derived normal distribution of the stationary *S*-allele frequency distribution (Eq. (7)) to the empirical *S*-allele frequency distribution. The empirical mean, 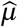, and variance, 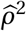, are used to fit a normal distribution. From this fitted normal distribution, we can directly compute the estimates for the number of *S*-alleles, 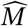, and the population size, 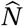, using Eq. (7):

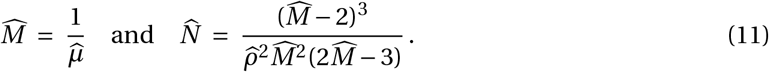

Note that by definition the estimated number of *S*-alleles, 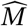, is exactly the observed number of *S*-alleles in the sample (the expectation of the allele frequency spectrum equals the number of observed *S*-alleles). This estimator therefore underestimates the true number of *S*-alleles in the population. Our first estimator, which takes into account the sampling process, corrects for this bias and therefore does not encounter this problem.

We also applied this second estimator to a simulated plant population of size 3,000 with different numbers of *S*-alleles. We find that the estimated population sizes mostly underestimate the true population size. This effect is most pronounced for small sample sizes (blue violin plots in Fig. **4B**). This is explained by smaller samples showing a larger variance in the allele frequency distribution, which translates to smaller values of the estimated population size 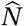 (Eq. (11)). The larger the sample size, the better is the population size approximated, yet there is still considerable variation in the estimates.

Lastly, we applied our estimator to an empirical dataset from Stoeckel et al. (2012), where a population of wild cherry (*Prunus avium*) was quasi-exhaustively sampled (*n* = 249). We estimated 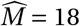 and 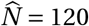, which is of the same order as, though underestimates, the census population size *n*. From the 95% confidence interval (CI) of the sample variance, we computed the 95% CI for the population size, which spans from 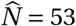 to 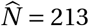, still not covering the sample size. The discrepancy between the estimation and the census population size can be due to a bias of our estimator. Another possible reason for this underestimation is violation of our modeling assumptions, *e.g*. spatial structure of the population or fecundity differences between individuals due to their size (Stoeckel et al., 2012). Overall, our results may serve as a starting point to develop new statistical estimators that can be used to infer ecological parameters like the effective population size from the diversity observed at a *S*-locus. In particular, as a perspective, it might be possible to combine Paxman’s model with a fitted normal distribution to jointly estimate the number of *S*-alleles and the population size, similar to our first attempt in SM, Section G.

## 4 Discussion

### Self-incompatibility as an historical illustration of probabilistic thinking

As emphasized by Wright (1937), one of the central goals of population genetics is to find the frequency distribution of genetic variants under various conditions, in particular, for convenience, under stationarity. GSI was one of the first genetic systems used to validate theoretical predictions from the young field of population genetics in the 1930-40’s. Before detailing how GSI was used to bring together theory and empirical observations, we first briefly summarize the general theory as stated by Fisher and Wright in the 1930’s (see Ishida and Rosales, 2020, for a detailed and general historical review).

The approach taken by Wright was to split the frequency changes into two parts: a deterministic part (describing mutation, migration or selection) and a stochastic part (describing random sampling errors). Wright generally first states how the frequency of a given allele *p_i_* changes in expectation from one generation to the next (by a difference equation), and second derives the form of the stochastic fluctuations, here denoted 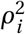. To derive the stochastic fluctuations, he usually assumes that the gene frequency dynamics are a succession of binomial draws, where the probability for a specific allele to be transmitted is equal to its frequency in the gamete offspring pool, assumed to be infinite. These allele frequencies in the offspring pool could have changed, compared to the parental generation, because of the processes of selection, migration or mutation (*e.g*. Wright, 1937). In this model, the stochastic fluctuations from one generation to the next are given by the variance of a binomial distribution 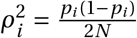. The deterministic change of the allele frequency Δ*p_i_* should appear in the binomial distribution parameters, but Wright explicitly assumes that it is negligible and thus that stochastic fluctuations only depend on the population size and the actual allele frequency, generally referred to as “genetic drift”.

Using a continuous approximation of the discrete stochastic process, Wright (1937, 1938) calculated the change of the mean and variance of the allele frequencies, giving a possible approximation of the distribution of allele frequencies for various cases regarding selection, migration and mutation, and its explicit stationary distribution. In his papers, Wright (1937, 1938) acknowledged that Fisher (1930) found similar results, despite Fisher’s approach being different. Thanks to a letter from the mathematician Kolmogorov, Wright (1945) realized that the form he had found for the allele distribution was the solution of the more general Fokker-Planck equation used in physics, which describes a stochastic process combining deterministic directed change on the one hand, and random motion on the other hand. At the same time, Malécot considered the change of the allele frequencies as a Markov process (reviewed in Nagylaki, 1989). In particular, Malécot rigorously demonstrated that assuming weak selection and a large population size, the form conjectured by Wright for the distribution of the allele frequencies was correct, probably the first application of the diffusion approximation to population genetics (Malécot, 1945). After the establishment of theoretical population genetics in the 1930s, it then remained to apply this general framework to a real scenario. GSI was an ideal problem since it combines selection, mutation and migration, with a parameter easily estimated from natural populations, the number of *S*-alleles.

Very rapidly Wright (1939) aimed at using the theory to resolve an apparent paradox from observations in one population of the flowering plant *O. organensis*. In this population, 45 self-incompatibility alleles were detected by Emerson (1938, 1939) even though the population size was thought to be not larger than one thousand individuals. Wright (1939) followed the general approach developed before (Fisher, 1930; Wright, 1938): he first provided the deterministic change of the frequency of a focal *S*-allele given the frequency of all the other *S*-alleles in the population, summarized in the parameter *R*, which measures the relative success of fertilization of the focal *S*-allele compared to the other *S*-alleles: 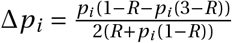. Wright remained elusive on the derivation and biological interpretation of *R*. We suspect that it can be seen as a selection coefficient, which depends on the genotype structure of the population.

Second, as in his previous works (Wright, 1937, 1938), Wright assumed that the stochastic fluctuations of the *i*-th *S*-allele frequency in a generation is given by the variance of a binomial distribution, from which he derived an approximation of the *S*-allele frequency distribution. As there was no explicit solution for the stationary distribution, Wright (1939) provided numerical results. He finally concluded that the large number of *S*-alleles observed in the *O. organensis* population would be possible if the mutation rate was as large as 10^−3^ – 10^−4^ per generation, or if the observed population was a small part of a much larger subdivided population with a low dispersal rate and a mutation rate of 10^−5^ – 10^−6^.

During almost twenty years, no important advances were made in the theory of stochastic processes applied to self-incompatibility systems. After experimental estimations of the mutation rate, found to be much lower than 10^-6^ in *O. organensis* by Emerson (1939) and Lewis (1948, 1949a), there was a revival of interest for having the best approximation of the frequency distribution of *S*-alleles in a population, launched in particular by a reexamination of the theory by Fisher (1958). Fisher (1958) challenged Wright’s (1939) model, called for developing the best stochastic approximation of the GSI system, and using it to identify the mechanisms underlying the pattern first highlighted by Emerson (1938). Fisher adopted the same approach as Wright (1939): first he derived the deterministic frequency change and then he assumed a form for the variance of the allelic change from a binomial distribution. In contrast to Wright however, Fisher (1958) motivated his deterministic change in allele frequency by an individual-based reasoning on the level of the plants. He was thus the first to give an explicit expression for the deterministic change of a given *S*-allele in the form given in Eq. (5) above. Moreover, he also proposed a different form, compared to Wright (1939), for the variance of the *S*-allele frequency change, 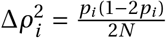. Similar to Wright, Fisher did not provide an explicit derivation of the infinitesimal variance from a probabilistic model. Finally, after deriving an allele frequency distribution under some technical simplification, Fisher came back to the plausible mechanisms explaining the large number of *S*-alleles observed in the *O. organensis* population (Emerson, 1938). He concluded that the population had most likely suffered from a strong population bottleneck, resulting in the population not being at drift-mutation equilibrium. Fisher discarded Wright’s (1939) hypothesis that the population could be strongly subdivided, with no clear reasons since Fisher did not consider any population structure or dispersal in his model. Fisher also took the opportunity to criticize that Wright (1939) only relied on numerical computations to analyze his model, and was not able to derive any explicit formula.

Wright (1960) answered Fisher’s (1958) criticisms mostly by comparing his 1939’s and new 1960’s models, and Fisher’s model through numerical simulations. Wright found no notable difference between the different models. Still he pointed out that Fisher’s approximation gave the worst estimation of *S*-allele frequency change among the three models. Overall, Wright (1960) and Fisher (1958), compared to Wright (1939), brought no important conceptual progress to the theory of GSI. Yet, these two papers illustrate that finding the correct form of approximation of the genetic dynamics of a GSI system was a major challenge for population genetics that led to another dispute between Wright and Fisher, beside the famous controversy about genetic dominance (Bagheri, 2006; Billiard and Castric, 2011).

A few years after this dispute, many papers were at least partly devoted to address the question of the correct approximation of the stationary distribution of *S*-allele frequencies in a finite population (Moran, 1962; Wright, 1964; Kimura and Crow, 1964; Ewens, 1964; Mayo, 1966; Ewens and Ewens, 1966; Wright, 1966). Yet again, no theoretical advances were made; most results relied on computer simulations that confirmed that Wright’s (1964) approximation was good enough. Later, Yokoyama and Nei (1979) and Yokoyama and Hetherington (1982) obtained new results by novel diffusion theory methods: instead of computing the allele frequency of a single allele, they computed the entire allele frequency distribution to directly infer the number of stably maintained *S*-alleles in a population. Yet again, similar to Wright’s different models (1939, 1960, 1964), they relied on a numerical approximation of the stationary distribution. Finally, and more recently, Muirhead and Wakeley (2009) derived recursion formulae to approximate the *S*-allele frequency distribution. This latest approach does not take a continuum limit (infinite population size limit) but approaches the problem in the discrete state space. While giving good results when compared to simulation results, this approach makes the formulae intractable and complicated to evaluate.

It is remarkable that the population genetics of self-incompatibility systems generated a lot of theoretical progress motivated by the aim to have theoretical predictions compatible with empirical observations. There was clearly a challenge for theoretical population geneticists, as if the stochastic theory of population genetics would be valuable only if it could help to explain the genetic diversity observed in natural populations, or at least if it could give correct predictions about parameter values like the mutation rate. In short, the main question that theoreticians tried to address was: what is the correct way to model the stochastic dynamics of allele frequencies in finite populations?

Two points of view opposed: the need for rigorous *vs*. practical derivations. Moran (1962) particularly criticized the lack of rigour of Wright’s approach, while Wright (1964) claimed that he only aimed for a *good* rather than a *justified* approximation. Such an opposition raised two difficulties: Is it possible to evaluate the validity of a model independently of data obtained from natural populations? How can the robustness and precision of the model’s approximation be evaluated? As emphasized by Moran (1962) and Malécot (reviewed in Nagylaki, 1989), these goals can be achieved by using a probabilistic way of thinking, *i.e*. by clearly justifying a stochastic model, and by properly deriving its approximation with probabilistic, mathematical tools. Here, we applied this theoretical framework to GSI.

### Reexamination of the number of *S*-alleles in a finite population

One aim of our study was to analytically revisit the results on the number of *S*-alleles under GSI, as derived by Wright (1939, 1964). Even though Wright’s results were numerically confirmed (see previous section), a theoretical confirmation was still lacking (to the best of our knowledge). Our results on the stationary distribution fit the numerically derived normal distribution obtained by Wright (1964) almost perfectly, *e.g*. Fig. 5, thus confirming Wright’s results also from a theoretical perspective.

**Figure 5:**
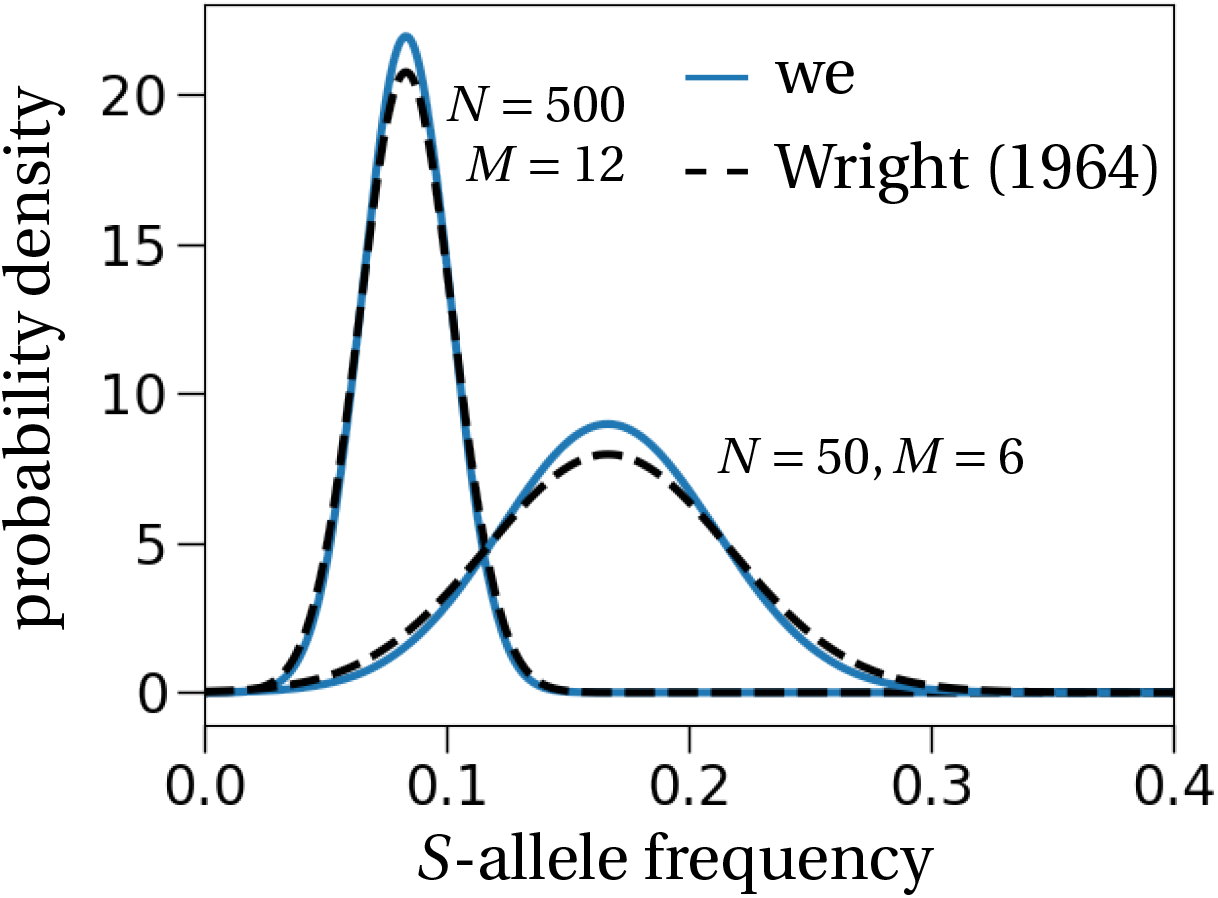
Stationary allele frequency distribution – comparison between Wright (1964) and our result. Our prediction is the result from Eq. (7) adapted to non-overlapping generations as considered by Wright (1964). The exact expression is stated in Eq. (C.9) in the SM. The black dashed lines correspond to the expressions computed by Wright (1964), for *M* = 12 *S*-alleles in a population of size *N* = 500 (left) and *M* = 6, *N* = 50 (right). We stress that Wright remains vague on the evaluation of the variance of this distribution and computes it numerically, *i.e*. he does not derive it analytically as we do (we refer to his discussion in section “Some examples” in Wright (1964)).

Following Wright’s (1939) strategy, we computed the number of *S*-alleles that can be maintained in a finite population of fixed size and for a fixed mutation rate (Fig. 2). Our analytical prediction slightly overestimates the simulated values. This is explained by the idealized assumption that all *S*-allele frequencies are centered around the steady state. Therefore, deviations from this stationary distribution, i.e., the rate of extinction of an *S*-allele is underestimated, which results in the overestimation of the simulated number of *S*-alleles. If we were to compute the number of *S*-alleles without the normal approximation of the stationary distribution, we would instead slightly underestimate the number of *S*-alleles as found in Czuppon and Rogers (2019) in the related haploid self-incompatibility (HSI) system, a mating system found for example in some fungi and ciliates.

Qualitatively, HSI is very similar to GSI because individuals can reproduce successfully only by mating with an individual with a different *S*-allele. This, as in GSI, produces negative frequency-dependent selection on the frequency of a focal *S*-allele, which results in a stable equilibrium frequency at an intermediate value. Recently, quantities like the number of mating types maintained in a population, the invasion probability, extinction dynamics, the effect of clonal reproduction and of fitness differences between different mating types have been addressed in this simpler genetic system (Constable and Kokko, 2018; Czuppon and Rogers, 2019; Czuppon and Constable, 2019; Krumbeck et al., 2020; Berríos-Caro et al., 2021). Even though simpler to analyze, we still suspect the qualitative results to carry over from the HSI to the GSI setting because the overall behavior of the two dynamical systems are similar.

For example, here our results were derived assuming neither fitness differences between different *S*-alleles, nor phenomena such as partial self-compatibility. Accounting for these mechanisms would alter the prediction of the number of *S*-alleles, e.g. it was shown that variation in zygote viability reduces the diversity at the *S*-locus (Uyenoyama, 2003), which is similar to results found in HSI systems (Krumbeck et al., 2020). In addition, we can speculate that the effect of self-fertilization on *S*-allele diversity maintained under GSI would be similar to that under HSI. We would therefore expect that the number of *S*-alleles is reduced for increasing rates of self-fertilization (Constable and Kokko, 2018; Czuppon and Constable, 2019; Berríos-Caro et al., 2021).

Considering a single panmictic population is another important simplification of our model. The subdivision of population and the migration rate between demes strongly affect the number of *S*-alleles maintained in a population (Wright, 1939; Schierup et al., 2000; Stoeckel et al., 2008). In particular, Schierup et al. (2000) showed that the total number of *S*-alleles at the scale of the whole metapopulation varies non-monotonically with the migration rate (the number of *S*-alleles is minimal for an intermediate value of the migration rate). This striking result was obtained with numerical simulations. A formal derivation of the *S*-diversity in a structured population was suggested by Wright (1939), yet surprisingly, no analytical progress has been made since then (to the best of our knowledge). We suspect that the framework developed in this manuscript could help to make analytical progress, *e.g*. by studying the edge cases of (fully) isolated populations, *i.e*., very low migration rates, and a well-mixed population without population structure, *i.e*., very high migration rates (essentially the model of this manuscript). Importantly, the invasion dynamics of new *S*-alleles into a local population will need to be taken into account.

### Invasion behavior under GSI and HSI

The invasion probability of a new *S*-allele under GSI, denoted by *φ* (Eq. (9)), can also be compared to the case of HSI. Czuppon and Rogers (2019) showed that a good approximation for the invasion probability of an *S*-allele in an HSI population with *M* resident *S*-alleles is 1/*M*. Here, for GSI, we find *φ > 1/M, i.e*. the invasion probability of a new *S*-allele is always larger for GSI than for HSI. This can be understood by comparing the ratio between the number of individuals that are compatible with the invading and with resident *S*-alleles in both situations. Applying the same reasoning that led to Eq. (10), we find

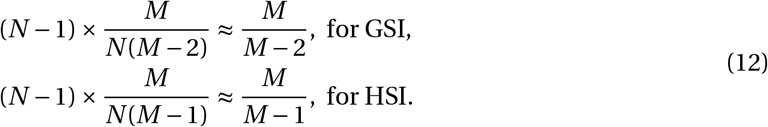

This shows that an invading (or rare) *S*-allele always has a larger fertilization advantage relative to a resident *S*-allele in GSI than in HSI. As a consequence, for similar population sizes, we would expect a higher *S*-allele diversity under GSI than under HSI dynamics because the rare allele advantage is stronger in the GSI mating system. The equations also show that as *M* increases, the difference between GSI and HSI vanishes as new *S*-alleles are getting closer to being neutral. Hence, as *M* increases, GSI and HSI tend to have similar invasion dynamics.

### *S*-haplotype diversification dynamics: does the underlying molecular architecture matter?

The number of *S*-alleles maintained in the population emerges from the balance between the *S*-allele loss rate by chance (genetic drift), the appearance of a new *S*-allele in the population (mutation rate), and its probability of invasion, which only depends on the number of resident *S*-alleles. As most stochastic models of GSI dynamics after Wright (1939), we assumed a one-step mutation model where mutations instantly generate new, fully functional *S*-alleles. If we further assume that mutations are rare enough so that the *S*-allele frequencies equilibrate between successive mutation events, we can quantify the diversification rate. That is, using (9) and a mutation rate *u*, we find that the diversification dynamics, i.e. the rate at which the number of *S*-alleles *n*(*t*) accumulates if the initial number of *S*-alleles is three, follows a square-root function of time: 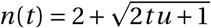 (mathematical details in Section H in the SM).

However, assuming a one-step mutation model is (very likely) an oversimplification as shown by the description of the genetic and molecular architectures underlying the SI phenotypes in different families (reviewed in Durand et al., 2020). Generally, a ‘superlocus’ composed of several genes is involved, which makes the appearance of new *S*-alleles more complicated as it requires several compensatory mutations allowing the self or non-self SI recognition. Using stochastic computer simulations, Gervais et al. (2011), Sakai (2016), Bod’ová et al. (2018) and Harkness et al. (2021) studied the diversification dynamics of *S*-alleles taking explicitly into account the succession of compensatory mutations invading a population. These multi-step mutation models were studied to better reflect the biological complexity associated with the appearance of a new *S*-allele. Gervais et al. (2011) and Bod’ová et al. (2018) showed that the diversification dynamics is due to complex interactions between the selection regimes of the different mutations present in the population (functional SI alleles or self-compatible alleles). Starting with a low number of *S*-alleles, they also showed that the diversification rate was first very fast, and rapidly reached either a plateau, where the number of *S*-alleles was stationary, or a regime where diversification rate was low.

Comparing the diversification dynamics expected under our one-step mutation model to the dynamics observed in the multi-step mutation models studied by Gervais et al. (2011), Sakai (2016), Bod’ová et al. (2018), and Harkness et al. (2021), our results suggest that the genetic architecture underlying SI seems to have only small effects on the diversification dynamics. Indeed, two types of diversification dynamics are observed in multi-step mutation models (Gervais et al., 2011; Sakai, 2016; Bod’ová et al., 2018; Harkness et al., 2021): either the number of *S*-alleles asymptotically approaches its stationary state, or the stationary number of *S*-alleles is rapidly reached after a diversification burst. These two diversification dynamics can be described by our one-step mutation model (Fig. 6), respectively when the mutation rate is low *vs*. high. When the mutation rate is low, the diversification dynamics follows a square-root function of time (Fig. **6A**). When the mutation rate is high, the diversification dynamics follows a square-root function of time truncated at the stationary number of *S*-alleles. The number of *S*-alleles accumulates and reaches the stationary phase quickly, which leads to an abrupt halt of increase in the number of *S*-alleles (Fig. **6B**). In stochastic simulations, the number of *S*-alleles then fluctuates around this stably maintained number of *S*-alleles. Overall, our results suggest that the diversification dynamics is mainly affected by an *effective mutation rate*, i.e., the rate of appearance of new *S*-alleles in multi-step models, rather than by the molecular and genetic details of the mechanism underlying SI. However, how this effective mutation rate is related to these mechanisms is an open issue that would need to be addressed by a specific model.

**Figure 6:**
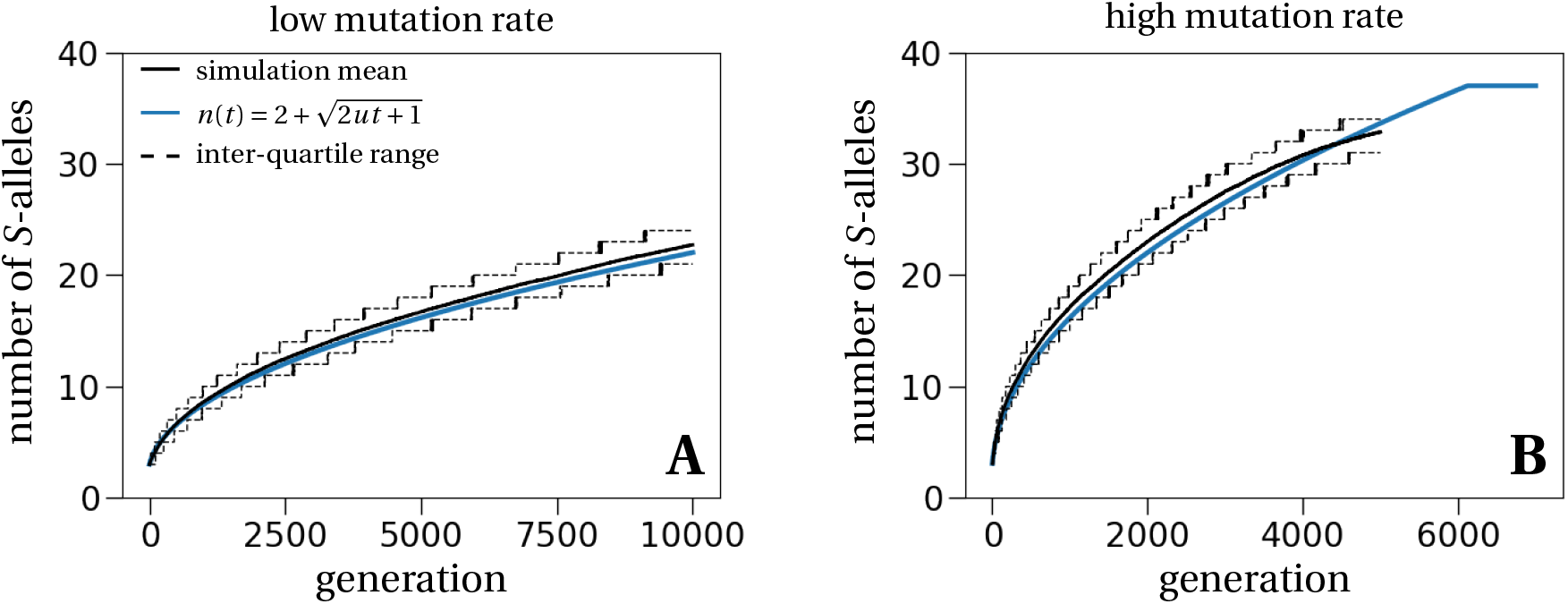
*S*-allele diversification dynamics in our one-step mutation model. The black solid line is the mean obtained from 1,000 stochastic simulations of a population with size *N* = 5,000 and started with three *S*-alleles in stationarity. The black dashed lines correspond to the 25- and 75-percentiles of the simulations. The blue line shows the theoretical prediction 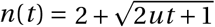, which fits well with the simulation results. The consistent underestimation is a result of the underestimation of the invasion probability (Fig. 3). The mutation rate is set to *u* = 1/50*N* in **A** and *u* = 1/10*N* in **B**. Simulations were stopped in **B** after 5,000 generations because of computational time constraints. The theoretically predicted number of stably maintained *S*-alleles for the high mutation rate is 37, which, as explained above, is slightly overestimating the number of stably maintained *S*-alleles in simulations.

The diversification dynamics derived in our one-step mutation model can also be used as a null model of *S*-allele diversification. One possible application could be to compare the distribution of times separating the appearance of *S*-alleles in phylogenies and to compare it to a square-root function of time. If a substantial discrepancy between empirical vs. null diversification dynamics is observed, this would suggest that, contrarily to what our model suggests, the molecular and genetic mechanisms underlying SI significantly affect the evolution of SI.

### Connecting theoretical results to data

Returning to the question about the observed diversity at the *S*-locus in *O. organensis*, Wright (1960) concluded that the empirically estimated low mutation rate obtained by Lewis (1948) would not be possible to maintain the 45 sampled *S*-alleles in an estimated population of 500 individuals. Instead, he proposed that population subdivision could be an explanation. However, as shown by simulations, population subdivision reduces the number of *S*-alleles in the population (Schierup, 1998), which contrasts the idea advocated by Wright (1939). Another hypothesis for the large number of observed *S*-alleles is a recent decrease in population size (Fisher, 1958) or that the population size has been underestimated (Levin et al., 1979). To our knowledge, the question about the large number of *S*-alleles in *O. organensis* has never been resolved, at least partly because appropriate theoretical and statistical tools were lacking. Dispersal kernels have been estimated in other GSI populations (*e.g*. Stoeckel et al., 2012), but estimating the effective population size is still necessary to identify and disentangle the processes underlying the observed diversity at the *S*-locus.

We used our approximation of the stationary distribution to derive a joint estimator for the number of *S*-alleles and the plant population size from a sample. The procedure is rather simple: we fit a normal distribution to the sampled allele frequency spectrum. This procedure will always estimate that there are as many *S*-alleles in the population as there are observed *S*-alleles in the sample. In this respect, our estimation is less accurate than Paxman’s (1963) estimator, which accounts for sampling error (details in SM, Section F). Yet, when estimating the population size from the allele frequency spectrum (by Eq. (11)), the moment estimator produces much better estimates than an extension of the method proposed by Paxman (1963) (compare Figs. **4A** and **B**). We therefore suggest to use the least-squares estimator, which accounts for the sampling process and is an extension of Paxman’s (1963) estimator, to estimate the number of *S*-alleles in the population and then to proceed with the moment estimator to compute the population size as defined in Eq. (11). The estimated population size is the idealized, in the sense that stationarity is assumed, population size that best explains the variance in the *S*-allele frequency distribution of the data sample. We recommend to take the estimator of population size with care because it has a tendency to underestimate the true population size (Fig. **4B**). One possible way to check if the obtained estimate is reasonable, is to compare the estimated population size with a prediction of the number of maintained *S*-alleles in the population (Fig. 2). If the population size is much too small compared to the number of observed *S*-alleles, the estimate is very likely an underestimate of the true population size. Similarly, if the estimated population size is too large compared to the predicted number of maintained *S*-alleles, e.g. by orders of magnitude, then the population size estimate should not be taken at face value. Alternatively, it is of course possible that other assumptions of our neutral model are violated, *e.g*. non-stationarity of the *S*-allele frequency distribution, which could suggest a recent population decline (or growth), or population structure, which means that the sampled population is part of a bigger metapopulation, which will affect the theoretical allele frequency distribution (in particular, the allelic frequency spectrum is expected not to be Gaussian anymore at intermediate migration rate: Wright, 1939; Schierup, 1998; Schierup et al., 2000; Muirhead, 2001; Stoeckel et al., 2008). A better understanding of the limitations of this new estimator is therefore needed to reliably apply it to data sets.

### Conclusion

To summarize, we have revisited the question of the possible number of *S*-alleles maintained in a finite population from a theoretical point of view. Our explicit expression of the stationary allele frequency distribution confirms the numerically obtained results by Wright (1964). Additionally, we provide an approximation for the invasion probability of a new *S*-allele and find that it is similar to the corresponding expression in populations with haploid self-incompatibility. Lastly, we define an estimator for the population size of a GSI population, which provides a starting point to assess the state of the sampled population. For example, a larger estimated diversity of *S*-alleles than predicted by the estimated population size in Fig. 2 might be indicative for a recent population decline. Lastly, our model might additionally be useful for conservation purposes. The reproductive output of plants can be reduced by pollen limitation, which may occur especially with low *S*-allele diversity in SI species (Leducq et al., 2010; Olivieri et al., 2016). Hence, if the population size is known, the prediction of the number of *S*-alleles for that size can help evaluate the extinction risk.

