## Supplementary Material for "Revisiting the number of self-incompatibility alleles in finite populations: from old models to new results"

#### Contents

**A Model definition and diffusion approximation** **S3**
A.1 Diffusion approximation of the genotype dynamics . . . . . S3
A.2 Diffusion approximation of the $S$-allele dynamics . . . . . S4

**B Stationary distributions** **S6**
B.1 Stationary distribution for the genotype frequency . . . . . S6
B.2 Stationary distribution for the $S$-allele frequency . . . . . S7
B.3 Comparison to simulation results . . . . . S8

**C Model with non-overlapping generations** **S9**
C.1 Genotype dynamics . . . . . S9
C.1.1 Deterministic dynamics . . . . . S9
C.1.2 Infinitesimal variance . . . . . S9
C.1.3 Stationary distribution . . . . . S10
C.2 Stationary distribution of the $S$-allele frequency . . . . . S10

**D Loss rate of an allele at frequency $1/2N$** **S11**

**E Invasion probability** **S12**
E.1 ... computed with the stochastic diffusion . . . . . S12
E.2 ... computed from the birth and death rates . . . . . S13
E.3 Comparing the different invasion probabilities . . . . . S13

**F Deriving the estimator from Paxman (1963)** **S15**

**G Estimating the number of $S$-alleles and the population size from a data sample** **S16**
G.1 Application to simulated data . . . . . S17

**H A null model for *S*-allele diversification** **S19**
H.1 Comparing the null model to simulations . . . . . . . . . . . . . . . . . . . . . . . . . . . . S19

#### A Model definition and diffusion approximation

We recapitulate the model from the main text. We consider a plant population of fixed size $N$ with overlapping generations. We assume that there is a fixed number of self-incompatibility alleles ($S$-alleles) in the population, and we denote this number by $M$. Individual plants are diploid and the number of plants with genotype $ij$ are denoted by $A_{ij}$. We do not order the alleles in this notation, i.e. $A_{ij} = A_{ji}$. An $A_{ij}$ individual can be fertilized by pollen of type $k \neq i, j$, i.e., we necessarily have $A_{ii} = 0$ for all $i$.

The detailed mechanisms that result in an increase or decrease of the number of individuals with a certain genotype are detailed in the main text. We just state the final result of the birth rate of genotype $ij$, denoted by $T^+_{ij}$, as given in Eq. (1) in the main text:

$$T^+_{ij} = \left( \frac{1}{2} \sum_{k\neq i,j} A_{jk} \frac{p_i}{1-p_j-p_k} + \frac{1}{2} \sum_{k\neq i,j} A_{ik} \frac{p_j}{1-p_i-p_k} \right) \frac{N-A_{ij}}{N}. \tag{A.1}$$

Applying the arguments from the main text, death processes (d-i), (d-ii) and (d-iii), the death rate $T^-_{ij}$ is computed as follows:

$$\begin{aligned} T^-_{ij} = &\Bigg( \underbrace{\frac{1}{2} \sum_{k\neq i,j} \sum_{l\neq k,i,j} A_{kl} \sum_{m\neq k,l} \frac{p_m}{1-p_k-p_l}}_{\text{death by event (d-i)}} \\ &+ \underbrace{\sum_{k\neq i} A_{ik} \sum_{m\neq i,k} \frac{p_m}{1-p_i-p_k} - \frac{1}{2} \sum_{k\neq i} A_{ik} \frac{p_j}{1-p_i-p_k}}_{\text{death by event (d-ii)}} \\ &+ \underbrace{\sum_{k\neq j} A_{jk} \sum_{m\neq j,k} \frac{p_m}{1-p_j-p_k} - \frac{1}{2} \sum_{k\neq j} A_{jk} \frac{p_i}{1-p_j-p_k}}_{\text{death by event (d-iii)}} \Bigg) \underbrace{\frac{A_{ij}}{N}}_{A_{ij}\text{ replacement}} \\ = &\left( \frac{1}{2} \sum_k \sum_{l\neq k} A_{kl} - \frac{1}{2} \sum_{k\neq i,j} A_{ik} \frac{p_j}{1-p_i-p_k} - \frac{1}{2} \sum_{k\neq j,i} A_{jk} \frac{p_i}{1-p_j-p_k} \right) \frac{A_{ij}}{N} \\ = &\left( N - \frac{1}{2} \sum_{k\neq i,j} A_{ik} \frac{p_j}{1-p_i-p_k} - \frac{1}{2} \sum_{k\neq j,i} A_{jk} \frac{p_i}{1-p_j-p_k} \right) \frac{A_{ij}}{N}. \end{aligned} \tag{A.2}$$

This is exactly Eq. (2) in the main text.

##### A.1 Diffusion approximation of the genotype dynamics

To obtain the diffusion approximation in a finite population of the gametophytic self-incompatibility model as stated above, we need to compute the infinitesimal mean and the infinitesimal variance (e.g. Czuppon and Traulsen, 2021; Otto and Day, 2007). Writing $a_{ij} = A_{ij}/N$ for the density of plants with genotype $ij$, the infinitesimal mean is given by

$$\begin{aligned}\frac{da_{ij}}{dt} &= \lim_{N\to\infty} \frac{T^+_{ij} - T^-_{ij}}{N} \\ &= \frac{1}{2}\left(\sum_{k\neq i,j} a_{jk}\frac{p_i}{1-p_j-p_k} + \sum_{k\neq i,j} a_{ik}\frac{p_j}{1-p_i-p_k}\right) - a_{ij} = \mu_{ij}\,.\end{aligned} \tag{A.3}$$

Analogously, the infinitesimal variance computes to

$$\begin{aligned}\sigma^2_{ij} &= \frac{T^+_{ij} + T^-_{ij}}{N^2} \\ &= \frac{1}{2N}\left(\sum_{k\neq i,j} a_{jk}\frac{p_i}{1-p_j-p_k} + \sum_{k\neq i,j} a_{ik}\frac{p_j}{1-p_i-p_k}\right) \\ &\quad + \frac{a_{ij}}{N}\left(1 - \sum_{k\neq i,j} a_{ik}\frac{p_j}{1-p_i-p_k} - \sum_{k\neq j,i} a_{jk}\frac{p_i}{1-p_j-p_k}\right).\end{aligned} \tag{A.4}$$

Even though we do not use the covariances between different genotypes in the diffusion approximation of the genotypes as stated in Eq. (3) in the main text, we still compute it here, because it will be useful to compute the infinitesimal variance of the $S$-allele dynamics. The infinitesimal covariance of genotypes $ij$ and $kl$ is given by

$$\begin{aligned}\gamma_{ij,kl} &= \mathbf{E}\left[(\Delta a_{ij} - \mathbf{E}[\Delta a_{ij}])(\Delta a_{kl} - \mathbf{E}[\Delta a_{kl}])\right] \\ &\approx -\frac{1}{2N}\left(\sum_{m\neq i,j}\left(a_{jm}\frac{p_i}{1-p_j-p_m} + a_{im}\frac{p_j}{1-p_i-p_m}\right)a_{kl}\right. \\ &\qquad \left. + \sum_{m\neq k,l}\left(a_{km}\frac{p_l}{1-p_k-p_m} + a_{lm}\frac{p_k}{1-p_l-p_m}\right)a_{ij}\right),\end{aligned} \tag{A.5}$$

where $\Delta a_{ij}$ denotes the infinitesimal change in $ij$-genotype frequency. The same methodology was used to find the diffusion approximation in the related haploid self-incompatibility system (Czuppon and Rogers, 2019).

#### A.2 Diffusion approximation of the $S$-allele dynamics

To find the diffusion approximation of the $S$-allele dynamics, we relate the $S$-allele frequency to the genotype frequencies. The transformation is given by

$$p_i = \sum_{j\neq i} a_{ij}/2\,. \tag{A.6}$$

Further noting that

$$\sum_{j\neq i}\sum_{k\neq i,j} a_{ik}\frac{p_j}{1-p_i-p_k} = \sum_{k\neq i}\sum_{j\neq i,k} a_{ik}\frac{p_j}{1-p_i-p_k} = \sum_{k\neq i} a_{ik} = 2p_i, \tag{A.7}$$

we obtain for the infinitesimal mean of the $S$-allele frequency change:

$$\mu_i = \frac{1}{2}\sum_{j\neq i}\mu_{ij} = \frac{p_i}{2}\left(\frac{1}{2}\sum_{j\neq i}\sum_{k\neq i,j} a_{jk}\frac{1}{1-p_j-p_k} - 1\right). \tag{A.8}$$

For the infinitesimal variance on the allele level, $\rho_i^2$, we make use of the following relation:

$$\rho_i^2 = \mathbf{Var}[\sum_{j\neq i} a_{ij}/2] = \frac{1}{4}\left(\sum_{j\neq i}\mathbf{Var}[a_{ij}] + \sum_{j\neq i}\sum_{k\neq i,j}\mathbf{Cov}[a_{ij},a_{ik}]\right). \tag{A.9}$$

We compute the terms one after the other. We start with the sum of the variances:

$$\begin{aligned}\sum_{j\neq i}\sigma_{ij}^2 &= \frac{1}{N}\left[\frac{1}{2}\left(\sum_{j\neq i}\sum_{m\neq i,j} a_{jm}\frac{p_i}{1-p_j-p_m} + 2p_i\right) + 2p_i - \sum_{j\neq i} a_{ij}\sum_{m\neq i,j} a_{im}\frac{p_j}{1-p_i-p_m}\right.\\ &\qquad\left. - \sum_{j\neq i} a_{ij}\sum_{m\neq j,i} a_{jm}\frac{p_i}{1-p_j-p_m}\right]\\ &= \frac{1}{N}\left[3p_i + \frac{1}{2}\sum_{j\neq i}\sum_{m\neq i,j} a_{jm}\frac{p_i}{1-p_j-p_m} - \sum_{j\neq i} a_{ij}\sum_{m\neq i,j} a_{im}\frac{p_j}{1-p_i-p_m}\right.\\ &\qquad\left. - \sum_{j\neq i} a_{ij}\sum_{m\neq i,j} a_{jm}\frac{p_i}{1-p_j-p_m}\right].\end{aligned} \tag{A.10}$$

For the sums of the covariances we have

$$\begin{aligned}\sum_{j\neq i}\sum_{k\neq i,j}\mathbf{Cov}[a_{ij},a_{ik}] &= -\frac{1}{2N}\sum_{j\neq i}\sum_{k\neq i,j}\left(\sum_{m\neq i,j}\left(a_{jm}\frac{p_i}{1-p_j-p_m} + a_{im}\frac{p_j}{1-p_i-p_m}\right)a_{ik}\right.\\ &\qquad\left. + \sum_{m\neq i,k}\left(a_{km}\frac{p_i}{1-p_k-p_m} + a_{im}\frac{p_k}{1-p_i-p_m}\right)a_{ij}\right)\\ &= -\frac{1}{2N}\sum_{k\neq i} a_{ik}\left(\sum_{j\neq i,k}\sum_{m\neq i,j} a_{jm}\frac{p_i}{1-p_j-p_m} + \sum_{m\neq i,k} a_{im}\right)\\ &\qquad - \frac{1}{N}p_i\left(\sum_{k\neq i,j}\sum_{m\neq i,k} a_{km}\frac{p_i}{1-p_k-p_m} + \sum_{m\neq i,j} a_{im}\right).\end{aligned} \tag{A.11}$$

Adding up Eqs. (A.10) and (A.11) and dividing by 4 (Eq. (A.9)) gives the infinitesimal variance of the allele dynamics, $\rho_i^2$.

### B Stationary distributions

We now compute an approximation for the stationary distribution of the genotype and $S$-allele frequencies in a population of size $N$ with $M$ different $S$-alleles. We know that in the deterministic steady state, which we abbreviate by $(*)$, the frequencies are, for all $i$, given by

$$(*) \qquad a^*_{ij} = \frac{2}{M(M-1)} \quad (j \neq i) \qquad \text{and} \qquad p^*_i = \frac{1}{M}. \tag{B.1}$$

This steady state is globally stable (Boucher, 1993). In this case we can compute the fluctuations around the steady state using a central limit theorem-type argument (Ethier and Kurtz, 1986; van Kampen, 2007; Czuppon and Traulsen, 2021). The same approach, in a slightly different notation, was applied in the related haploid self-incompatibility system in Czuppon and Constable (2019) to derive the number of mating types in a finite population under facultative clonal reproduction.

#### B.1 Stationary distribution for the genotype frequency

We first compute the stationary distribution for the genotypes for a given number of $S$-alleles $M$. The fluctuations around the steady state, denoted by $U$, are defined by

$$U_t = \frac{1}{\sqrt{N}}\left(a^N_{ij}(t) - a_{ij}(t)\right), \tag{B.2}$$

where $a^N_{ij}(t)$ are the frequency dynamics of genotype $ij$ in the finite population size system, which is indicated by the superscript $N$. The deterministic dynamics as defined in Eq. (A.3) are described by $a_{ij}(t)$. The stochastic process $U_t$ then describes the error between the finite population size model and the deterministic trajectories of the genotype frequency dynamics, scaled by $\sqrt{N}$, reminiscent of the central limit theorem.

From Eq. (B.2), we find that the fluctuation process $U_t$ satisfies the following stochastic differential equation (we refer to Czuppon and Traulsen (2021), Section 5.2, for more details):

$$dU_t = \frac{1}{\sqrt{N}}\left(\frac{\partial \mu_{ij}}{\partial a_{ij}} U dt\right) + \sqrt{\sigma^2_{ij}}\, dW_t. \tag{B.3}$$

Note that we have neglected the covariance terms and derivatives of $\mu_{ij}$ with respect to $a_{kl}$ $(kl \neq ij)$ in Eq. (3) in the main text. This introduces an error in the approximation of the fluctuations for low numbers of $S$-alleles $M$, i.e., when these neglected terms are largest. Based on the comparison of the theoretical stationary distribution with stochastic simulation results, we conclude that this error is negligible (Fig. A(a)). For small genotype frequencies, corresponding to large values of $M$, the diffusion approximation does not appropriately describe the dynamics at the extinction boundary (Assaf and Meerson, 2017), i.e., the stationary distribution is not well described by a Gaussian distribution.

Evaluated at the deterministic steady state $(*)$, the fluctuation process $U_t$ becomes an Ornstein-Uhlenbeck process with variance $-\sigma^2_{ij}/(2\mu'_{ij})$, with all terms evaluated at $(*)$. The derivative of the deterministic dynamics in stationarity simplifies to:

$$\left.\frac{\partial \mu_{ij}}{\partial a_{ij}}\right|_* = \begin{cases} -\frac{3}{2}, & M = 3, \\ -1, & M \geq 4. \end{cases} \tag{B.4}$$

For $M = 3$ we have used that $a_{23} = 1 - a_{12} - a_{13}$, for $M \geq 4$ we can always choose the ordering of the genotypes so that $a_{ij}$ only appears once in Eq. (A.3) and $\sum_{i=1}^M \sum_{j>1} a_{ij} = 1$ is still satisfied. The variance and covariance in stationarity reduce to

$$\left.\sigma^2_{ij}\right|_* = \frac{4(M-2)(M+1)}{NM^2(M-1)^2} \quad \text{and} \quad \left.\gamma_{ij,kl}\right|_* = -\frac{8}{NM^2(M-1)^2}. \tag{B.5}$$

Putting things together, we find that the stationary distribution of genotypes is then given by

$$\psi_{ij}^* \sim \begin{cases} \mathcal{N}\left(\frac{2}{M(M-1)}, \frac{4(M-2)(M+1)}{3NM^2(M-1)^2}\right), & M=3, \\ \mathcal{N}\left(\frac{2}{M(M-1)}, \frac{2(M-2)(M+1)}{NM^2(M-1)^2}\right), & M \geq 4, \end{cases} \tag{B.6}$$

where $\mathcal{N}(\mu, \sigma^2)$ denotes a normal distribution with mean $\mu$ and variance $\sigma^2$.

#### B.2 Stationary distribution for the $S$-allele frequency

We follow the same approach as above, i.e., we compute the stationary variance by applying the central limit theorem for density-dependent Markov processes (Ethier and Kurtz, 1986): we define a fluctuations process $V_t$ analogously to Eq. (B.2), which then satisfies the analogous dynamics as obtained in Eq. (B.3):

$$dV_t = \frac{1}{\sqrt{N}}\left(\frac{\partial \mu_i}{\partial p_i} V dt\right) + \sqrt{\rho_i^2} dW_t. \tag{B.7}$$

Again, we will evaluate the process at the steady state $(*)$. Similarly to the genotype frequencies, due to our constraint of a fixed population size, we need to write the frequency $p_M$ in terms of the other $S$-allele frequencies: $p_M = 1 - \sum_{l=1}^{M-1} p_l$. The computation of the derivative of the term $\mu_i$ is a bit more involved compared to the computation on the genotype level because variation in $p_i$ is due to variation in the plant frequencies $a_{i\bullet}$. Therefore, we approximate the derivative $\partial\mu_i/\partial p_i$ as follows:

$$\frac{\partial \mu_i}{\partial p_i} \approx \sum_{j \neq i} \frac{\partial \mu_i}{\partial a_{ij}}. \tag{B.8}$$

Recalling that $p_i = \sum_{j\neq i} a_{ij}/2$, this computes to

$$\begin{aligned}\frac{\partial \mu_i}{\partial a_{il}} = &\underbrace{\frac{1}{4}\left(\frac{1}{2}\sum_{j\neq i}\sum_{k\neq i,j} a_{jk}\frac{1}{1-p_j-p_k} - 1\right)}_{\text{contribution from the factor } p_i/2} + \underbrace{\frac{p_i}{2}\left(\frac{1}{2}(-1)\frac{1}{1-p_M-p_{M-1}}\times 2\right)}_{\text{contribution from } a_{M(M-1)} = 1-\sum_{(j,k)\neq(M,M-1)} a_{jk}} \\ &+ \underbrace{\frac{p_i}{2}\left(\frac{1}{2}\sum_{j\neq i,l,M} a_{jl}\frac{1}{(1-p_j-p_l)^2}\times\frac{1}{2}\times 2\right)}_{\text{contribution from } a_{il} \text{ in the term } p_l \text{ in the denominator}} + \underbrace{\frac{p_i}{2}\left(\frac{1}{2}\sum_{j\neq i,l,M} a_{jM}\frac{1}{(1-p_j-p_M)^2}\times(-1)\times 2\right)}_{\text{contribution from } p_M = 1-\sum_{j\neq M} p_j \text{ in the denominator}} \\ &+ \underbrace{\frac{p_i}{2}\left(\frac{1}{2}a_{lM}\frac{1}{(1-p_l-p_M)^2}\times\left(-\frac{1}{2}\right)\times 2\right)}_{\text{contribution from } a_{lM} \text{ denominator}}.\end{aligned} \tag{B.9}$$

Applying the stationarity condition $(*)$, this simplifies to

$$\begin{aligned}\frac{\partial \mu_i}{\partial p_i} &\approx \sum_{j\neq i}\frac{\partial \mu_i}{\partial a_{ij}} \\ &= (M-1)\times\left(0 - \frac{1}{2(M-2)} + \frac{M(M-3)}{2(M-2)^2(M-1)} - \frac{(M-3)}{(M-1)(M-2)^2} - \frac{1}{2(M-1)(M-2)^2}\right) \\ &= -\frac{2M-3}{2(M-2)^2}\end{aligned} \tag{B.10}$$

The variance $\rho_i^2$ in stationarity is given by

$$\rho_i^2\Big|_* = \frac{1}{4}(M-1)\sigma_{ij}^2\Big|_* + \frac{1}{4}(M-1)(M-2)\gamma_{ij,kl}\Big|_* = \frac{M-2}{M^2}. \tag{B.11}$$

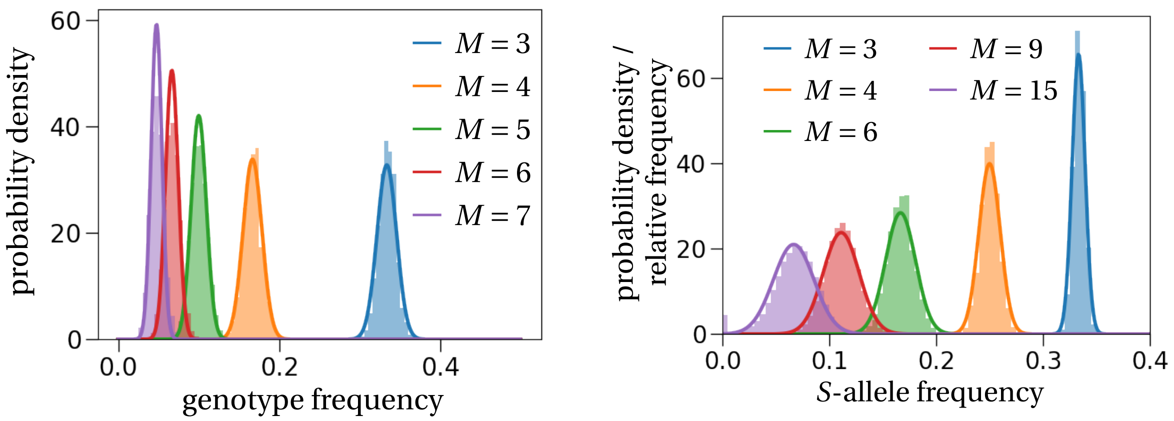


(a) genotype distribution

(b) allele distribution

Figure A: **Stationary distributions of genotypes and *S*-alleles.** As predicted by the theory (solid lines computed by (a) Eq. (B.6) and (b) Eq. (B.13)), the stationary distribution of the genotype and allele frequencies are normal distributions centered at (a) $2/M(M-1)$ and (b) $1/M$. The histograms in both panels are obtained from 100 stochastic simulations that ran for 1,200 generations in a population of size $N = 1,000$, where we recorded the frequency of the same $S$-allele every generation. To ensure no dependence on the initial state we started recording frequencies of a single genotype or allele after 200 generations. If the number of $S$-alleles becomes too large, e.g. $M = 15$, alleles might get lost, which is shown by the purple bar at the frequency equal to zero.

Gathering all the terms, we arrive at the stationary variance of the Ornstein-Uhlenbeck process:

$$-\frac{\rho_i^2|_{x^*}}{2N}\left(\frac{\partial \mu_i}{\partial p_i}\bigg|_*\right)^{-1} \approx \frac{(M-2)^3}{NM^2(2M-3)}. \tag{B.12}$$

The stationary allele distribution is then given by

$$\psi \sim \mathcal{N}\left(\frac{1}{M}, \frac{(M-2)^3}{NM^2(2M-3)}\right). \tag{B.13}$$

This explains Eq. (7) in the main text.

##### B.3 Comparison to simulation results

We compare both stationary distributions to simulations results. We observe that as long as the numerical value of the steady state $(*)$ is sufficiently far away from the extinction boundary, the analytical approximations give a reasonably good fit, even for low numbers of $S$-alleles, where the neglected covariance terms would have the most effect (Fig. A).

### C Model with non-overlapping generations

With the rates of birth and death as identified in the Moran model of GSI we know that the total reproduction rate of the population is (Eq. (A.1) without the death term)

$$\frac{1}{2}\sum_{i=1}^{M}\sum_{j\neq i} T_{ij}^{+} = \frac{1}{2}\sum_{i=1}^{M}\sum_{j\neq i}\frac{1}{2}\left(\sum_{k\neq i,j} a_{jk}\frac{p_i}{1-p_j-p_k} + \sum_{k\neq i,j} a_{ik}\frac{p_j}{1-p_i-p_k}\right) = 1\,. \tag{C.1}$$

Further, in a model with non-overlapping generations, the probability of a certain genotype configuration in the next generation is a multinomial distribution. The probabilities of drawing a $ij$-genotype are given by

$$\pi_{ij} = \frac{1}{2}\sum_{k\neq i,j} a_{jk}\frac{p_i}{1-p_j-p_k} + \frac{1}{2}\sum_{k\neq i,j} a_{ik}\frac{p_j}{1-p_i-p_k}\,. \tag{C.2}$$

#### C.1 Genotype dynamics

##### *C.1.1 Deterministic dynamics*

We denote by $A^t$ the genotype configuration in generation $t$ and by $A_{ij}^t$ the number of $ij$-genotype plants in generation $t$. The mean dynamics on the genotype level for a single genotype can then be derived by computing (Etheridge, 2012; Czuppon and Traulsen, 2021)

$$\mathbf{E}\left[A_{ij}^{t+1} - A_{ij}^{t}\middle|A^t\right] = N\pi_{ij} - A_{ij}^t\,. \tag{C.3}$$

Transforming to densities, $a_{ij}^t = A_{ij}^t/N$ and rescaling time by $1/N$ we find

$$\begin{aligned}\mathbf{E}\left[\frac{A_{ij}^{t+1/N} - A_{ij}^{t}}{N}\middle|A^t\right] &= \pi_{ij} - a_{ij}^t \\ &= \frac{1}{2}\sum_{k\neq i,j} a_{jk}\frac{p_i}{1-p_j-p_k} + \frac{1}{2}\sum_{k\neq i,j} a_{ik}\frac{p_j}{1-p_i-p_k} - a_{ij} = \mu_{ij}\,,\end{aligned} \tag{C.4}$$

where $\mu_{ij}$ is given by Eq. (4) in the main text.

##### *C.1.2 Infinitesimal variance*

To describe the stochastic fluctuations in a finite population, we compute the infinitesimal variance given by (time steps in $1/N$)

$$\begin{aligned}N\mathbf{E}\left[(a_{ij}^{t+1/N} - a_{ij}^t)^2|A^t\right] &= \frac{1}{N}\mathbf{E}\left[(A_{ij}^{t+1/N} - A_{ij}^t)^2|A^t\right] = \frac{1}{N}\left(\mathbf{Var}\left[A_{ij}^{t+1/N}|A^t\right] + \left(\mathbf{E}\left[A_{ij}^{t+1/N}|A^t\right] - A_{ij}^t\right)^2\right) \\ &= \frac{1}{N^2}\left(N\pi_{ij}(1-\pi_{ij}) + \mu_{ij}^2\right) \\ &= \frac{1}{2N}\left(\sum_{k\neq i,j} a_{jk}\frac{p_i}{1-p_j-p_k} + \sum_{k\neq i,j} a_{ik}\frac{p_j}{1-p_i-p_k}\right) \\ &\quad\left(1 - \frac{1}{2}\left(\sum_{k\neq i,j} a_{jk}\frac{p_i}{1-p_j-p_k} - \sum_{k\neq i,j} a_{ik}\frac{p_j}{1-p_i-p_k}\right)\right) + O\left(\frac{1}{N^2}\right) \\ &= \sigma_{ij}^2\,,\end{aligned} \tag{C.5}$$

where we used the large $O$-notation for terms of order $1/N^2$ or smaller.

The covariance terms can be computed analogously:

$$\begin{aligned}
\gamma_{ij,kl} &= \mathbf{E}\left[(\Delta a_{ij} - \mathbf{E}[\Delta a_{ij}])(\Delta a_{kl} - \mathbf{E}[\Delta a_{kl}])\right] \\
&= -\frac{N}{N^2}\pi_{ij}\pi_{kl} \\
&= -\frac{1}{4N}\left(\sum_{u\neq i,j} a_{ju}\frac{p_i}{1-p_j-p_u} + \sum_{u\neq i,j} a_{iu}\frac{p_j}{1-p_i-p_u}\right) \\
&\quad \times \left(\sum_{u\neq k,l} a_{ku}\frac{p_l}{1-p_k-p_u} + \sum_{u\neq k,l} a_{lu}\frac{p_k}{1-p_l-p_u}\right).
\end{aligned} \tag{C.6}$$

##### C.1.3 Stationary distribution

We can compute the stationary distribution of genotypes by exactly the same procedure as in the Moran-type model. Since the deterministic drift is exactly the same as in the Moran-type model, the deterministic steady state $a^*, p^*$ is the same as in Eq. (B.1). The same holds for the partial derivative of $\mu_{ij}$ with respect to $a_{ij}$, which is the same as stated in Eq. (B.4). For the stationary variance we then find

$$\sigma^*_{ij} = \frac{2(M-2)(M+1)}{NM^2(M-1)^2}, \tag{C.7}$$

which is exactly half of the variance of the Moran-type model (compare to Eq. (B.5)). This is a known difference between Moran- and Wright-Fisher models that arises due to the different sampling schemes of non-overlapping and overlapping generations (Czuppon and Traulsen, 2021).

Analogously, the covariance in stationarity reduces to

$$\gamma^*_{ij,kl} = -\frac{4}{NM^2(M-1)^2}, \tag{C.8}$$

where again there is a two-fold difference to the Moran-type model (Eq. (B.5)).

#### C.2 Stationary distribution of the $S$-allele frequency

As we have seen in the previous section, the variance (and covariance) in stationarity is exactly half of the respective quantities of the Moran-type model. Therefore, following the same steps as in Section B, the stationary distribution of $S$-alleles in the Wright-Fisher updating scheme is given by

$$p^* \sim \mathcal{N}\left(\frac{1}{M}, \frac{(M-2)^3}{2NM^2(2M-3)}\right). \tag{C.9}$$

#### D Loss rate of an allele at frequency $1/2N$

We compute the death rate of a plant that is the last remaining individual with a certain $S$-allele at frequency $1/2N$ in the Moran-type model. Say it is allele $i$ that is at frequency $1/2N$ and the last remaining plant with this allele is an $ij$-plant. From the death rate in Eq. (A.2), we find

$$\begin{aligned}\frac{T_{ij}^{-}}{N} &= \left(1-\frac{1}{2}\sum_{k\neq i,j} a_{ik}\frac{p_j}{1-p_i-p_k}-\frac{1}{2}\sum_{k\neq j,i} a_{jk}\frac{p_i}{1-p_j-p_k}\right)a_{ij} \\ &\approx \left(1-0-\frac{1}{2}(M-2)\frac{2}{(M-1)(M-2)}\frac{1/2N}{1-2/(M-1)}\right)\frac{1}{N} \\ &\approx \left(1-\frac{1}{2N(M-3)}\right)\frac{1}{N} \\ &\approx \frac{1}{N}.\end{aligned} \tag{D.1}$$

### E Invasion probability

#### E.1 ... computed with the stochastic diffusion

We apply diffusion theory to compute the survival probability of a novel $S$-allele when initially rare (e.g. Otto and Day, 2007; Czuppon and Traulsen, 2021). To this end, we need to compute the scale function of the stochastic diffusion, defined by

$$S(x) = \int^x \exp\left(-2\int^y \frac{\mu(z)}{\rho^2(z)}dz\right)dy, \tag{E.1}$$

with the functions $\mu(z)$ and $\rho(z)$ as defined in Eqs. (A.8) and (A.9).

Assuming that there are $M$ resident alleles in the population with frequencies close to the equilibrium $(*)$, we obtain

$$\mu(x) \approx \frac{x}{2}\left(\frac{1}{2}M(M-1)\frac{2}{M(M-1)}\frac{1}{1-2/M}-1\right) = \frac{x}{M-2}. \tag{E.2}$$

If we further assume that we can neglect terms of order $x^2$, a reasonable assumption as long as the novel $S$-allele frequency is very low, we find

$$\begin{aligned}\rho^2(x) &\approx \frac{1}{2N}\left(\frac{x}{2}\left(1+\frac{1}{2}M(M-1)\frac{2}{M(M-1)}\frac{1}{1-2/M}\right)+x\left(1-2x-M(M-1)\frac{2}{M(M-1)}\frac{x}{1-2/M}\right)\right)\\ &\approx \frac{x}{2N}\left(\frac{M-1}{M-2}+1\right)\\ &= \frac{x}{2N}\left(\frac{2M-3}{M-2}\right).\end{aligned} \tag{E.3}$$

The scale function $S(x)$ then simplifies to

$$S(x) = \frac{2M-3}{4N}\left(1-\exp\left(-\frac{4N}{(2M-3)}x\right)\right) \tag{E.4}$$

The survival probability (or hitting probability of the frequency $1/(M+1)$), denoted by $\varphi$, is then given by (e.g. Czuppon and Traulsen, 2021)

$$\varphi = \frac{S(1/2N)-S(0)}{S(1/(M+1))-S(0)} = \frac{1-\exp\left(-\frac{2}{(2M-3)}\right)}{1-\exp\left(-\frac{4N}{(2M-3)(M+1)}\right)} \approx 1-\exp\left(-\frac{2}{(2M-3)}\right), \tag{E.5}$$

where we have used that $N \gg M^2$ in the last approximation. Lastly, assuming that $M$ is large enough, so that terms of order $1/M^2$ are negligible, we find the survival probability

$$\varphi \approx \frac{2}{2M-3}. \tag{E.6}$$

**Remark** *We find the same expressions as in Eqs.* (E.2) *and* (E.3) *if we vary the resident allele and genotype frequencies in response to the increasing new allele frequency $x$ under the same assumptions as above ($x^2 \ll 1$). Specifically, for $i \leq M$ we write*

$$p_i = \frac{1-x}{M}, \tag{E.7}$$

*where $x$ is the allele frequency of the invading allele. This equation can be obtained by solving $Mp_i + x = 1$. A similar argument, $\frac{M(M-1)}{2}a_{ij} + xc = 1$, where $c$ needs to be determined by the condition $\frac{2}{M(M+1)} = \frac{2(1-cx)}{M(M-1)}$, results in*

$$a_{ij} = \frac{2(1-cx)}{M(M-1)} \quad \textit{with} \quad c = \frac{M+3}{2}, \tag{E.8}$$

*for $i, j \leq M$.*

#### E.2 ... computed from the birth and death rates

Alternatively, one can compute the invasion probability of a novel $S$-allele directly by the individual-based birth and death rates. In this case, the establishment probability $\varphi_{\text{bd}}$ is given by

$$\varphi_{\text{bd}} = 1 - \frac{\sum_{\alpha=1}^{\infty} \frac{T_1^- \cdots T_\alpha^-}{T_1^+ \cdots T_\alpha^+}}{1 + \sum_{\alpha=1}^{\infty} \frac{T_1^- \cdots T_\alpha^-}{T_1^+ \cdots T_\alpha^+}}, \tag{E.9}$$

where $T_\alpha^{+/-}$ are the transition rates to increase or decrease the number of alleles in the population from $\alpha$ to $\alpha+1$ or $\alpha-1$, respectively.

To compute these fractions of allele transition rates we need to transform the genotype transition rates from Eqs. (1) and (2) from the main text to the allele level. We computed these rates already when we derived the deterministic dynamics of a focal $S$-allele in Eq. (5) from the main text. Assuming that the $i$−th allele is rare and that the other $M$ $S$-alleles are close to the $M$-equilibrium we find

$$\frac{1}{2}\sum_{j\neq i}\sum_{k\neq i,j} a_{jk}\frac{1}{1-p_j-p_k} \approx \frac{1}{2}M(M-1)\frac{2}{M(M-1)}\frac{1}{1-2/M} = \frac{M}{M-2}. \tag{E.10}$$

We identify this value as the fraction $T_\alpha^+/T_\alpha^-$ and thus obtain

$$\varphi_{\text{bd}} = 1 - \frac{\sum_{\alpha=1}^{\infty}\left(\frac{M-2}{M}\right)^\alpha}{1+\sum_{\alpha=1}^{\infty}\left(\frac{M-2}{M}\right)^\alpha} = \frac{2}{M}. \tag{E.11}$$

This value is twice as large as the invasion probability in a haploid self-incompatibility system, where the invasion probability is $1/M$ (Czuppon and Rogers, 2019).

#### E.3 Comparing the different invasion probabilities

We compare the three different estimates from Eqs. (E.5), (E.6) and (E.11) to simulation results. Surprisingly, the best fit is produced by Eq. (E.6), which is an approximation for large numbers of resident alleles $M$ of Eq. (E.5). Yet, even for low values of the number resident alleles $M$, it better describes the data than the non-approximated result (blue vs. orange lines in Fig. B). Unfortunately, we have no explanation for this. The estimate obtained from the birth and death rates (Eq. (E.11); green line in Fig. B) consistently overestimates the simulated invasion probabilities, except for $M = 3$. This is explained by the assumption that the rare allele advantage diminishes with increasing frequency of the invading allele, whereas this approximation assumes it to be constant and equal to the value at frequency $1/2N$, i.e., the largest value of the rare allele advantage.

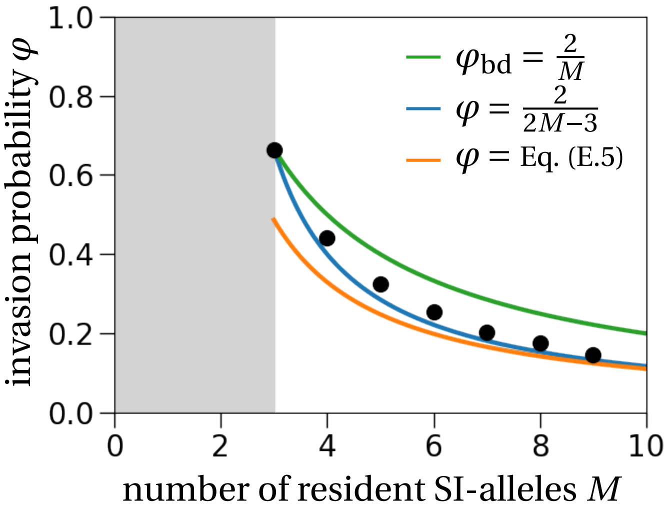


Figure B: **Comparison of the different estimates of the invasion probability.** The approximation of the invasion probability obtained from the stochastic diffusion (blue line, Eq. (E.6)) best fits the simulations (black dots). Interestingly, for low values of resident alleles $M$, it fits the simulation results better than the non-approximated result in Eq. (E.5) (orange line), which underestimates the simulated values and just describes the invasion probability well for larger $M$, where it coincides with the approximation (blue line). The invasion probability computed from the birth-death rates (green line, Eq. (E.11)) overestimates the simulation results, except for $M = 3$, where it takes the same value as the invasion probability from Eq. (E.6) (blue line).

### F Deriving the estimator from Paxman (1963)

The probability distribution of drawing a specific $S$-allele from the total population is a binomial distribution; this is analogous to drawing a single color from an urn with differently colored balls. This means that in a sample of plants of size $n$, the possible number of sampled $S$-alleles is between 1 and $n$ (one cannot not draw an allele, which is why 0 is not a possible value in this reasoning).

The probabilities can then be derived as follows. For a focal $S$-allele to be observed only once in the whole sample, the probability is given by

$$\mathbf{P}(1) = \binom{n}{1} 2\,\frac{1}{M}\left(1-\frac{1}{M}\right)^{n-1}\left(1-\frac{1}{M-1}\right)^{n-1}. \tag{F.1}$$

The binomial term is the number of possibilities for this allele to be distributed over the $n$ sampled plants. The term $2/M$, where $M$ is the number of $S$-alleles that we want to estimate, gives the probability of this allele to be observed in a plant (2 because of diploidy). The terms with the exponent $n-1$ describe the probability that the remaining $n-1$ plants do not carry the focal $S$-allele, which happens with probability $(1-1/M)(1-1/(M-1))$. Generalizing this idea, we find for the probability that a focal $S$-allele is observed $k$ times in a sample by

$$\mathbf{P}(k) = \binom{n}{k}\left(\frac{2}{M}\right)^{k}\left(\left(1-\frac{1}{M}\right)\left(1-\frac{1}{M-1}\right)\right)^{n-k} \approx \binom{n}{k}\left(\frac{2}{M}\right)^{k}\left(1-\frac{2}{M}\right)^{n-k}. \tag{F.2}$$

This binomial distribution is biased in the sense that one cannot observe the value 0, which means that we need to condition this distribution to positive values. This introduces a correction factor $1-(1-2/M)^n$.

To estimate the number of $S$-alleles from a sample of size $n$, we now equate the mean of this binomial distribution with the observed average. The average number of observations of a randomly chosen $S$-allele in the data is $2n/\widehat{M}$, where $\widehat{M}$ is the number of observed $S$-alleles in the sample. We obtain

$$\frac{2n}{M(1-(1-2/M)^n)} = \frac{2n}{\widehat{M}} \iff \widehat{M} = M\left(1-\left(1-\frac{2}{M}\right)^{n}\right), \tag{F.3}$$

which is the estimate derived in Paxman (1963).

#### G Estimating the number of $S$-alleles and the population size from a data sample

Instead of comparing the allele frequency distribution to a normal distribution as done in the main text, we here use the same reasoning as applied in Paxman (1963) to derive a joint estimate for the number of $S$-alleles in a population and the population size. To this end, we will, as in SM, Section F, make use of the analogy with an urn model, where balls with different colors are drawn from an urn. The different colors correspond to the different $S$-alleles and the number of balls with a specific color is normally distributed according to the stationary distribution.

We denote by $N_i$ the random variable of the number of balls corresponding to the $S$-allele of type $i$, so that $N_i \sim \mathcal{N}(2N/M, 4N^2\sigma^2_{p^*})$, where $\sigma^2_{p^*}$ is the variance of the stationary distribution of $S$-alleles (Eq. (B.11)). We denote by $n$ the plant sample size, so that $2n$ is the number of sampled $S$-alleles. Let $\widehat{M}$ be the observed number of $S$-alleles from the sample. Similarly to Paxman (1963), one can show that the number of observations of a focal $S$-allele is distributed according to a truncated binomial distributed with success probability $N_i/N = x_i$ and truncation at 0 (it is not possible to make no observation of an $S$-allele, so that the binomial distribution needs to be conditioned to be positive). The mathematical details are given in SM, Section F above. This truncation introduces a factor $(1-(1-x_i)^n)^{-1}$. We denote by $Y$ the random variable corresponding to the number of observations of a focal $S$-allele, say type $i$. Then $Y$ is a compound random variable, because the probability distribution is given by

$$\mathbf{P}(Y=k|N_i) = \binom{n}{k} x_i^k (1-x_i)^{n-k} \left(1-(1-x_i)^n\right)^{-1}, \tag{G.1}$$

which depends on the random variable $N_i$ through $x_i$. Therefore, the expectation of this compound random variable is given by (using the law of total expectation): $\mathbf{E}[Y] = \mathbf{E}_{N_i}[\mathbf{E}_Y[Y|N_i]]$. The index next to the expectation indicates the random variable to which this expectation corresponds to. Setting $\mathbf{E}[Y] = 2n/\widehat{M}$, which is the average number of observations of an $S$-allele in the sample, we obtain an extension of the maximum likelihood estimator defined by Paxman (1963).

Putting things together, we obtain

$$\mathbf{E}[Y] = \int_0^1 f(x) n x \frac{1}{1-(1-x)^n} dx \overset{!}{=} \frac{2n}{\widehat{M}}, \tag{G.2}$$

where $f(x)$ is the probability distribution of a normal distribution corresponding to the random variable $N_i/N$. Note that replacing $x$ by its mean $2/M$, and ignoring the integral in that case (formally we use $f(x)dx = \delta_{\{2/M\}}(x)$ with $\delta_z$ being the Dirac measure), recovers the estimate from Paxman (1963) (details in SM, Section F). Since we have two unknowns, the population size $N$ and the number of $S$-alleles $M$, we need a second equation to determine these values. To this end, we use the variance of the random variable $Y$ that we equate with the variance obtained from the data. The general equation that needs to be solved reads:

$$\mathbf{V}[Y] = \mathbf{E}_{N_i}[\mathbf{V}_Y[Y|N_i]] + \mathbf{V}_{N_i}[\mathbf{E}_Y[Y|N_i]] \overset{!}{=} \frac{1}{n}\sum_{i=1}^{\widehat{M}} \left(y_i - \frac{2n}{\widehat{M}}\right)^2, \tag{G.3}$$

where $y_i$ corresponds to the abundance of the $i$-th $S$-allele in the data.

Writing out the terms explicitly, we find

$$\mathbf{E}_{N_i}[\mathbf{V}[Y|N_i]] = \int_0^1 f(x)\left(\frac{nx(1-x)}{1-(1-x)^n} - \frac{n^2x^2(1-x)^n}{(1-(1-x)^n)^2}\right) dx, \tag{G.4}$$

where we used

$$\begin{aligned}\mathbf{V}[Y|N_i=x] &= \mathbf{E}[Y^2|N_i=x]-\mathbf{E}[Y|N_i=x]^2\\ &= \left(nx(1-x)+n^2x^2\right)\frac{1}{1-(1-x)^n}-n^2x^2\frac{1}{(1-(1-x)^n)^2}\\ &= nx(1-x)\frac{1}{1-(1-x)^n}-n^2x^2\frac{(1-x)^n}{(1-(1-x)^n)^2}\,.\end{aligned} \tag{G.5}$$

The second term computes to:

$$\begin{aligned}\mathbf{V}_{N_i}[\mathbf{E}_Y[X|N_i]] &= \mathbf{E}_{N_i}[\mathbf{E}_Y[Y|N_i]^2]-\underbrace{\mathbf{E}_{N_i}[\mathbf{E}_Y[Y|N_i]]^2}_{=(\mathbf{E}[Y])^2}\\ &= \int_0^1 f(x)n^2x^2\frac{1}{(1-(1-x)^n)^2}dx-\left(\int_0^1 f(x)nx\frac{1}{1-(1-x)^n}dx\right)^2\,.\end{aligned} \tag{G.6}$$

#### G.1 Application to simulated data

Based on Eqs. (G.2) and (G.3), we defined two least square estimators. The first one is unweighted and is given by

$$\mathrm{LSQ(data)}=\left(\mathbf{E}[Y]-\frac{2n}{\widehat{M}}\right)^2+\left(\mathbf{V}[Y]-\frac{1}{n}\sum_{i=1}^{\widehat{M}}\left(y_i-\frac{2n}{\widehat{M}}\right)^2\right)^2\,, \tag{G.7}$$

where $\mathbf{E}[Y]$ and $\mathbf{V}[Y]$ are given by the left-hand sides of Eq. (G.2) and Eq. (G.3), respectively.

The second estimator is weighted so that the mean and the variance contribute equally to the parameter estimation. In the unweighted version, the mean of the $S$-allele frequency distribution has a larger effect on the parameter estimate than the variance because its absolute value is larger. The weighted version of the estimator defined in Eq. (G.7) is

$$\mathrm{LSQ}_w\mathrm{(data)}=\frac{\left(\mathbf{E}[Y]-\frac{2n}{\widehat{M}}\right)^2}{2n/\widehat{M}}+\frac{\left(\mathbf{V}[Y]-\frac{1}{n}\sum_{i=1}^{\widehat{M}}\left(y_i-\frac{2n}{\widehat{M}}\right)^2\right)^2}{\frac{1}{n}\sum_{i=1}^{\widehat{M}}\left(y_i-\frac{2n}{\widehat{M}}\right)^2}\,. \tag{G.8}$$

We applied both estimates to the same simulated data set as already used in the main text (Fig. 4 in the main text). The estimate of the number of $S$-alleles is good for both least-square estimators, except for cases where the numerical optimizer does not converge. To estimate the number of $S$-alleles and the population size from a data sample, we minimize the least-square estimators above. Unfortunately, the estimators exhibit a plateau along the population size axis (Fig. C), which causes problems during the numerical optimization. This results in unsuccessful optimizations and the minimization yields unreasonable results.

The estimates for the population sizes vary widely for both least-square estimates (Fig. D). Importantly, the estimates are further away from the true population size (dashed line in Fig. D) than the ones obtained by the moment-estimator in the main text (Fig. 4B in the main text). As already mentioned, the optimization does not always converge. For the unweighted least-square estimator (Eq. (G.7)), we have: for $M = 15$ with 50, 100, 200 sampled plants out of ten times the procedure converged 4, 4, 6 times; with $M = 20$ and 50, 100, 200 sampled plants it converged in 3, 8, 8 cases; with $M = 25$ and 50, 100, 200 sampled plants it converged in 3, 9, 10 cases. The weighted least-square estimator (Eq. (G.8)) showed the following convergence behavior: for $M = 15$ with 50, 100, 200 sampled plants out of ten times the procedure converged 9, 6, 7 times; with $M = 20$ and 50, 100, 200 sampled plants it converged in 7, 3, 3 cases; with $M = 25$ and 50, 100, 200 sampled plants it converged 6, 8, 10 times.

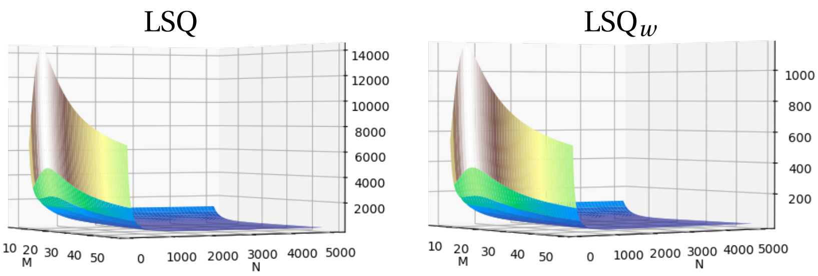


Figure C: **Least-squares estimators for the joint estimation of the number of $S$-alleles and the population size.** Both least-square estimators (left: unweighted from Eq. (G.7); right: weighted from Eq. (G.8)) are very flat in the dimension of the population size, which makes this parameter very hard to estimate. Parameters: true $M = 20$, true $N = 3000$, sample size = 100.

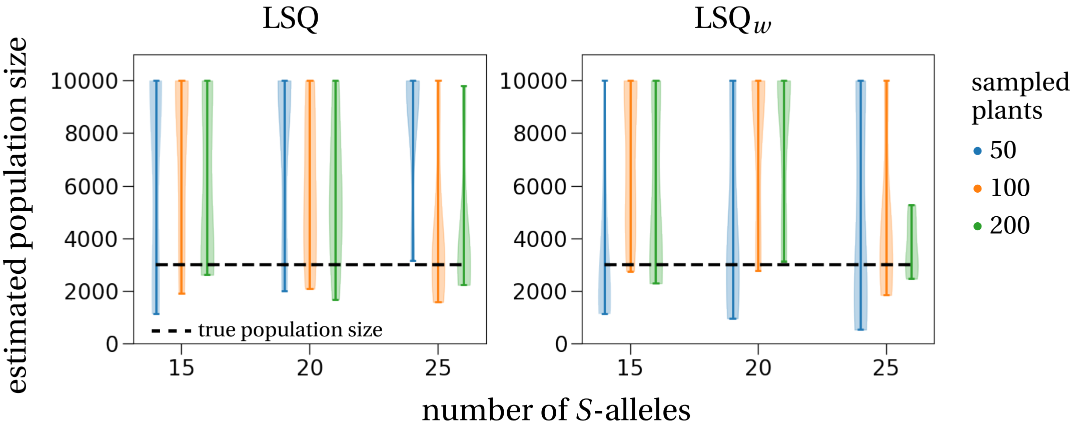


Figure D: **Estimated population sizes from simulated data.** The estimated population sizes vary widely for both least-square estimators (left: unweighted – Eq. (G.7); right: weighted – Eq. (G.8)) and tend to overestimate the true population size (3000 plants, dashed line). The simulated population of size 3000 with 15, 20 or 25 different $S$-alleles was started close to its steady state. The random sample is taken after 500 generations. The violinplots show the distribution of estimated population sizes from the 10 different samples taken from the same data set, implemented by a hypergeometric distribution (sampling without replacement). Blue violinplots correspond to 50 sampled plants, orange violinplots to 100 sampled plants, and green violinplots to 200 sampled plants. The estimates are truncated at a maximal estimated population size of 10000. Increasing the sample size seems to increase the chance for successful convergence, which is indicated by the broader parts of the violinplots being closer to the true population size (dashed lines).

Overall, due to the lack of stable convergence and the worse estimates of the population sizes when compared to the less sophisticated moment estimator from the main text, i.e., fitting a normal distribution to the allele frequency spectrum, we do not recommend to use the least-square estimators that we defined above to be applied to data.

### H A null model for $S$-allele diversification

In this section, we derive the diversification rate of $S$-alleles. To study diversification, we initialize a population of a fixed size $N$ with three different $S$-alleles at its equilibrium. New $S$-alleles arise with a mutation rate $u$, i.e., the waiting time for a new mutation to appear is $1/u$. Each new mutation has a probability of $\varphi(M)$ to establish, where $\varphi(M)$ is the invasion probability from Eq. (E.6), where the current number of $S$-alleles in the population is $M$. Thus, on average the new $S$-allele needs $1/\varphi(M)$ invasion attempts to succeed. The time to reach $n$ different $S$-alleles can then be computed as follows:

$$T(n) = \frac{1}{u}\sum_{k=3}^{n-1}\varphi(k) = \frac{1}{u}\sum_{k=3}^{n-1}\frac{2k-3}{2} = \frac{1}{2u}\left(n(n-1)-6-3(n-3)\right) = \frac{n^2-4n+3}{2u} = \frac{(n-2)^2-1}{2u}. \quad \text{(H.1)}$$

Inverting this function gives the number of $S$-alleles over time:

$$M(t) = \sqrt{2ut+1}+2, \quad \text{(H.2)}$$

which is the expression stated in the discussion in the main text and referred to as the null model of diversification.

#### H.1 Comparing the null model to simulations

We compare the theoretical prediction to the mean number of $S$-alleles from 1,000 stochastic simulations. Each simulation was started with three $S$-alleles and in stationarity, i.e., genotype frequencies were all at $1/3$. The population size was set to $N = 5,000$, which can stably maintain around 33 $S$-alleles for a low mutation rate ($u = 1/(50N)$) and 37 $S$-alleles for a high mutation rate ($u = 1/(10N)$), based on our prediction from the main text. The mutation rates are set to $u = 1/(50N)$ and $u = 1/(10N)$ per unit time step. The mutation rates per generation therefore are $u = 1/50$ and $u = 1/10$.

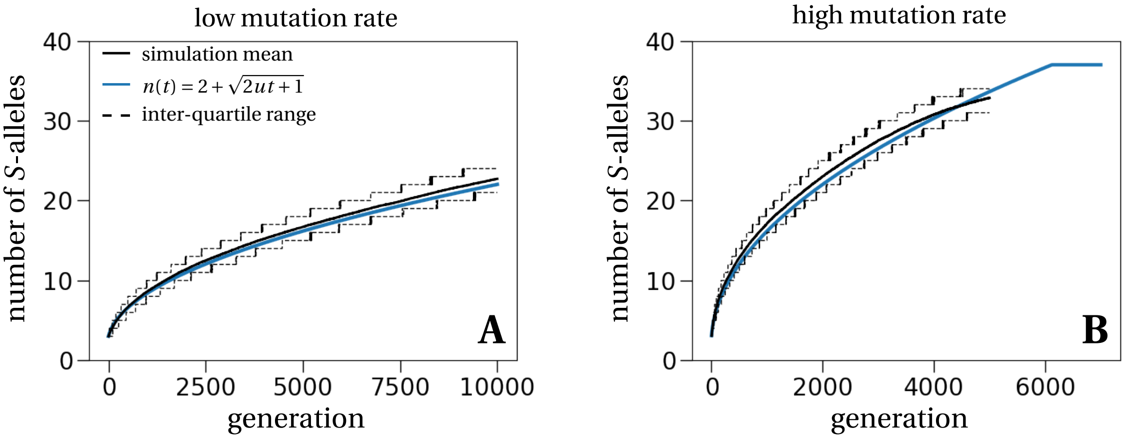


Figure E: **Diversification dynamics of $S$-alleles.** (same as Fig. 6 in the main text) The black solid line is the mean obtained from 1,000 stochastic simulations started with three $S$-alleles and in stationarity. The black dashed lines correspond to the 25- and 75-percentiles of the simulations. The blue line shows the theoretical prediction $n(t) = 2+\sqrt{2ut+1}$, which fits well with the simulation results. The consistent underestimation is a result of the underestimation of the invasion probability (Fig. B). The mutation rate is set to $u = 1/50N$ in **A** and $u = 1/10N$ in **B**.
